# USP7 Inhibition Promotes Early Osseointegration in Senile Osteoporotic Mice

**DOI:** 10.1101/2023.11.15.567232

**Authors:** F Zhou, Z Wang, H Li, D Wang, Z Wu, F Bai, Q Wang, W Luo, G Zhang, Y Xiong, Y Wu

**Affiliations:** State Key Laboratory of Oral Diseases, National Clinical Research Center for Oral Diseases, West China Hospital of Stomatology, Sichuan University, Chengdu, China; Department of Oral Implantology, West China Hospital of Stomatology, Sichuan University, Chengdu, China

**Author notes:** Correspondence should be addressed to: Yingying Wu, D.D.S, Ph.D. State Key Laboratory of Oral Diseases, National Clinical Research Center for Oral Diseases, and Department of Oral Implantology, West China Hospital of Stomatology, Sichuan University, 14 Third Section, Renmin Nan Road, Chengdu 610041, China.

**Keywords:** senile osteoporosis, osseointegration, USP7, efferocytosis, senolysis

## Abstract

Although elderly osteoporotic patients have similar implant survival rates compared to those of normal individuals, they require longer healing periods to achieve proper osseointegration. This may be related to chronic inflammatory responses and impaired stem cell repair functions in the osteoporotic bone microenvironment. Recently, the deubiquitinating enzyme, ubiquitin-specific peptidase 7 (USP7), was found to regulate macrophage immune response and modulate stem cell osteogenic differentiation. The selective inhibitor of USP7, P5091, has also been found to promote bone repair and homeostasis in osteoporotic conditions. However, the roles of USP7 and P5091 in osteoimmunology and dental implant osseointegration under senile osteoporotic conditions remain unclear. In this study, USP7 depletion and P5091 were showed to inhibit inflammation in senescent bone marrow derived macrophages (BMDMs) and promote osteogenic differentiation in aged BMSCs. Furthermore, mRNA-Seq revealed that USP7 depletion could enhance efferocytosis in senescent BMDMs through the EPSIN1/ low-density lipoprotein receptor-related protein 1 (LRP1) pathway and selectively induce apoptosis (senolysis) in aged BMSCs. In senile osteoporotic mice, we found that the osseointegration period was prolonged compared to young mice, and P5091 promoted the early stage of osseointegration, which may be related to macrophage efferocytosis around the implant. Collectively, this study suggests that USP7 inhibition may accelerate the osseointegration process in senile osteoporotic conditions by promoting macrophage efferocytosis and aged BMSCs apoptosis. This has implications for understanding the cellular interactions and signaling mechanisms in the peri-implant bone microenvironment under osteoporotic conditions. It may also provide clinical significance in developing new therapies to enhance osseointegration quality and shorten the edentulous period in elderly osteoporotic patients.

## INTRODUCTION

Senile osteoporosis, an age-related bone metabolic disease, leads to decreased bone quality, impaired bone healing, alveolar bone loss and tooth loss (Nicopoulou-Karayianni et al. 2009). Osteoporotic patients exhibit poorer primary stability and more pronounced marginal bone loss (Lemos et al. 2023; Merheb et al. 2016). Consequently, osteoporotic patients often require longer time to achieve proper osseointegration (Tsolaki et al. 2009). Simultaneously, antiresorptive drugs such as bisphosphonates may increase the risk of medication-related osteonecrosis of the jaw (MRONJ) which is triggered by implant surgery (Jung et al. 2023; Sher et al. 2021). Therefore, researching new therapies to enhance osseointegration in senile osteoporotic patients has considerable clinical significance.

In a tibial implant model of aged mice, studies have found an excessive innate immune response in the peri-implant tissues of aged mice (Turajane et al. 2021). Overactive innate immune response in aged individuals can generate a chronic inflammatory microenvironment, which is called inflammaging (Ferrucci and Fabbri 2018). In aged bone tissue, inflammaging is exacerbated by senescent myeloid cells and osteocytes, which exhibit the most pronounced senescence-associated secretory phenotype (SASP), and produce pro-inflammatory cytokines (Farr et al. 2016). In recent years, a novel strategy called senolysis, which selectively induces apoptosis in senescent cells and reduces SASP, has been proposed. P5091, a selective inhibitor targeting ubiquitin-specific peptidase 7 (USP7), could function as a senolytic drug to eliminate senescent fibroblasts in mice (He et al. 2020). Therefore, USP7, a key member of deubiquitinating enzymes (DUBs) regulating the ubiquitin-proteasome system (UPS), may play a vital role in senile osteoporotic microenvironment.

UPS is the primary protein degradation system, serving various functions such as clearing misfolded proteins, regulating cellular stress and signaling pathways, and controlling DNA damage repair (Park et al 2020). While this process is intricately regulated by the deubiquitination system. DUBs can cleave the ubiquitin chains from substrate proteins, thereby sparing them from degradation (Komander et al. 2009). USP7, a well-studied member of DUBs, was found to regulate cell differentiation, cell senescence and inflammatory response in age-related bone diseases (Shen et al. 2023). USP7 maintains self-renewal capacity of BMSCs via NANOG and SOX2 (Kim et al. 2022). However, Ji et al. argued that USP7 inhibited Wnt/β-catenin signaling via stabilizing axin (Ji et al. 2019). Similar contradictory results are also observed in animal disease models. Recently, Lin et al. found that P5091 could inhibit osteoclast differentiation to alleviate bone loss in osteoporotic mice (Lin et al. 2023). On the contrary, Xie et al. found that USP7 inhibited osteoclast differentiation by stimulating STING, thereby reducing the level of bone loss (Xie et al. 2023). The different function of USP7 may be attributed to different animal models. Lin et al. used OVX-induced osteoporosis model while Xie et al. used RANKL-induced skull bone loss model. It indicated that the function of USP7 varied among different disease model. Therefore, USP7 may be an important therapeutic target for bone-related diseases. However, whether USP7 inhibition promotes implant osseointegration or not still needs further investigations.

In this study, we found that USP7 inhibition could selectively eliminated aged BMSCs and enhance the efferocytosis of senescent BMDMs via EPSIN1/low-density lipoprotein receptor-related protein 1 (LRP1) pathway. Through systemic injection of P5091, we found that USP7 inhibition could improve early stage of osseointegration in senile osteoporotic mice via senolysis and efferocytosis. Collectively, USP7 inhibition enhanced BMDMs efferocytosis and increase aged BMSCs senolysis, which efficiently clear apoptotic senescent cell and inhibit macrophage inflammaging to promote early osseointegration.

## Materials and Methods

Materials and methods were described in Appendix Materials and Methods. Animal experiments were conducted in accordance with the ARRIVE 2.0 guidelines.

## Results

### Osseointegration was delayed in senile osteoporotic mice accompanied by elevated USP7 expression

Based on microCT results, 18-month-old mice exhibited a significant osteoporotic phenotype in femurs and maxilla (Fig. S1). We used 6-month-old mice and 18-month-old mice as the young group and the aged group, and implanted titanium implants anterior to the first maxillary molar. The bone volume around implants in aged mice consistently remained lower than that in young mice. However, both aged and young mice eventually achieved similar levels of bone-implant contact (BIC), while aged mice took 6 weeks to reach the BIC level that young mice achieved at 4-week. (Fig. 1A). These results indicated that aged mice had a slower rate of osseointegration. The differences in peri-implant bone volume and osseointegration level were most pronounced at 2-week and 4-week. USP7 expression around the implants was overall lower in young mice compared to aged mice. In aged mice, USP7 expression was significantly higher at 2-week, and became lower at 4-week (Fig. 1B-E). Further, we found that USP7 and p16INK4a were minimally expressed in young BMSCs and BMDMs, whereas in aged BMSCs and senescent BMDMs, both USP7 and p16INK4a were highly expressed in the cytoplasm and nucleus (Fig. 1F-I; Fig. S2). This suggested that USP7 might play a significant role in the early stage of osseointegration in aged mice.

**Fig. 1.**
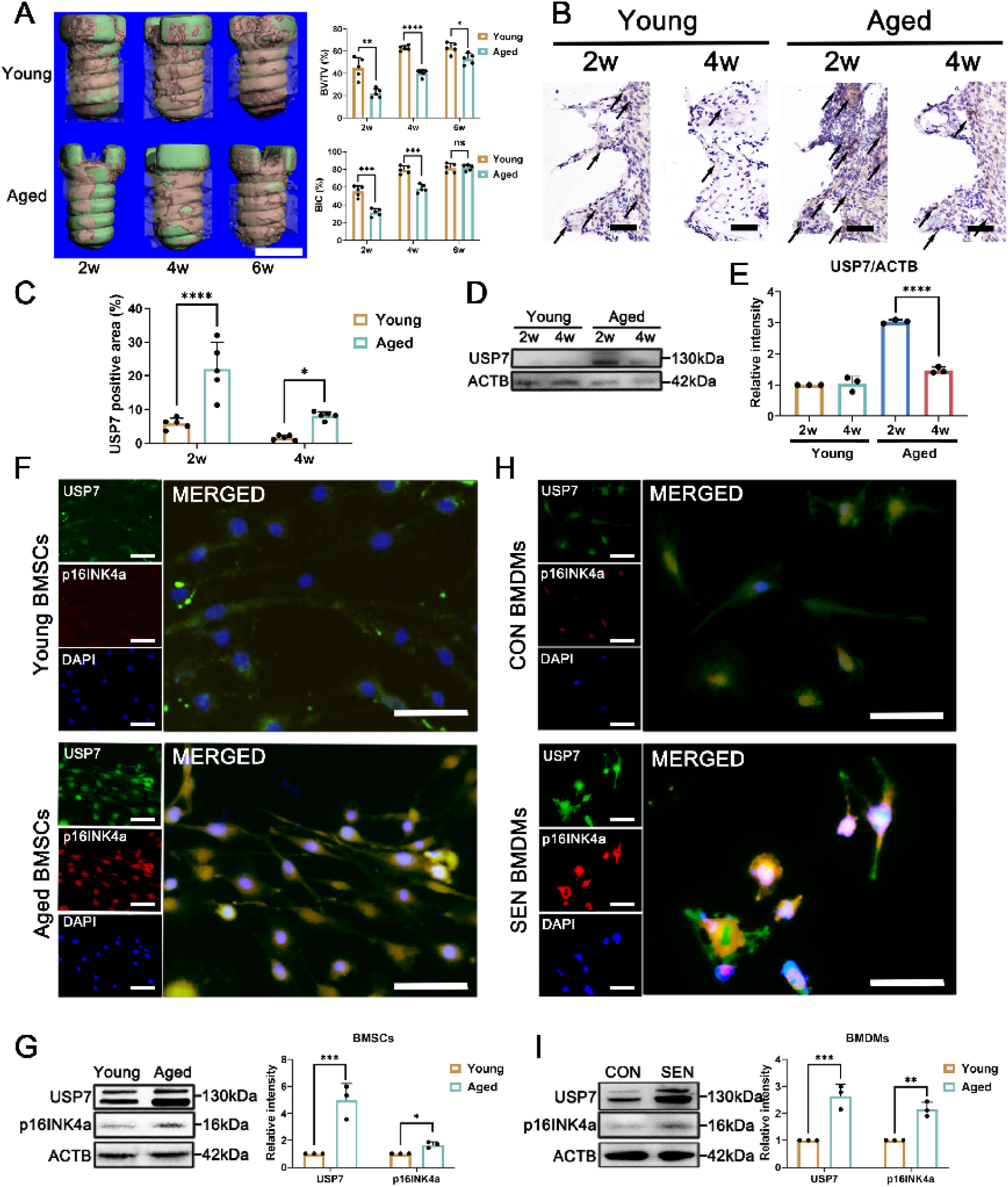
Senile osteoporotic mice exhibited a delayed osseointegration and higher expression of USP7. (A-B) Three-dimensional reconstruction image of alveolar bone around implants after 2 weeks and 4 weeks, scale bar, 600µm. Quantitative MicroCT analysis was conducted at the 200µm region around the implants, n = 5. (B) Immunohistochemical staining of USP7 around the implant, the black arrows indicated USP7-positive cells, scale bar, 100µm. (C) Quantification of USP7-positive area in bone tissue around implants, n = 5. (D-E) Western blot analysis and quantification of USP7 expression in the bone tissue around implants, n = 3. (F) Immunofluorescence staining of USP7, p16INK4a, and DAPI in young and aged BMSCs. (G) Western blot analysis and quantification of USP7 and p16INK4a expression in the BMSCs, n = (H) Immunofluorescence staining of USP7, p16INK4a, and DAPI in control and senescent BMDMs. (I) Western blot analysis and quantification of USP7 and p16INK4a expression in the BMDMs, n = 3. BV/TV, bone volume per tissue volume; Tb.N, trabecular number; Tb.Th, trabecular thickness; Tb.Sp, trabecular separation; BIC, bone-implant intact; Young, young BMSCs; Aged, aged BMSCs; CON, control BMDMs; SEN, senescent BMDMs . Data are shown as the mean ± SD; ns, not significant, \**P* < 0.05, \*\**P* < 0.01, \*\*\**P* < 0.001, \*\*\*\**P* < 0.0001.

### USP7 inhibition ameliorated inflammaging via NLRP3 pathway in senescent BMDMs

Subsequently, we used CM from aged BMSCs to induce senescent BMDMs and conducted mRNA-Seq (Fig. 2A). USP7 was depleted by shRNA targeting USP7 (Fig. S3A-B). USP7 depletion did not alter cell viability of BMDMs (Fig. S3C). KEGG pathway enrichment analysis revealed that in USP7-depleted senescent BMDMs, the TNF-α, NFκB, Toll-like signaling pathway, and Nod-like receptor pathways were downregulated (Fig. 2B). GSEA also showed that NLRP3 pathway and innate immune response was significantly inhibited by USP7 depletion (Fig. 2C). WB results showed that in senescent BMDMs, the protein level of p16INK4a was increased, accompanied by elevated levels of cleaved CASP1, GSDMD-NT, and cleaved IL-1β proteins. USP7 depletion significantly inhibited the expression of these proteins (Fig. 2D-E). Besides, we used P5091 to inhibit USP7. First, we found that 5µm of P5091 did not show significant cytotoxicity to BMDMs and BMSCs (Fig. S4A-B). Subsequently, we found that USP7 depletion and P5091 both significantly downregulated the mRNA levels and secretion of inflammatory cytokines such as IL-1β, IL-6, and TNF-α in senescent BMDMs (Fig. S3D-E; Fig. S4C-D).

**Fig. 2.**
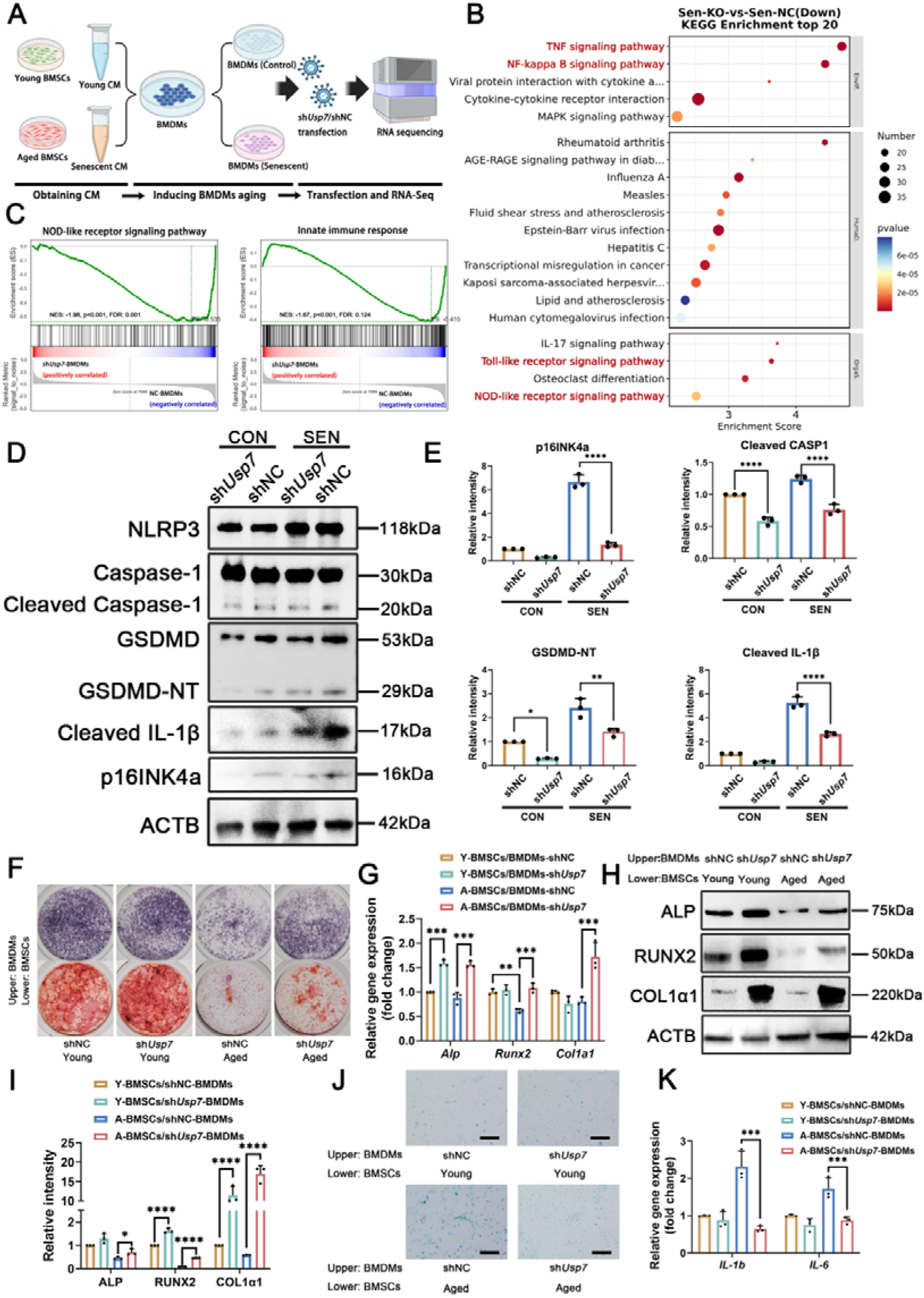
USP7 depletion inhibited inflammaging in senescent BMDMs and promoted osteogenesis of aged BMSCs co-cultured with BMDMs. (A) schematic diagram showing the induction of senescent BMDMs and mRNA-Seq analysis. (B) KEGG pathway enrichment analysis of senescent and USP7-depleted senescent BMDMs. (C) GSEA showing decreased enrichment of innate immune response pathway-associated genes and Nod-like receptor (NLR) signaling pathway-associated genes in USP7-depleted senescent BMDMs. (D) Western blot analysis and (E) quantification of p16INK4a, cleaved caspase-1, GSDMD-NT, and cleaved IL-1β in different groups of BMDMs, n = 3. (F) Representative images of ALP and ARS staining in young/aged BMSCs co-cultured with different groups of BMDMs for 7d and 14d. (G) qPCR analyses of the mRNA levels of *Alp*, *Runx2*, *Col1a1* in young/aged BMSCs co-cultured with BMDMs for 7d, n = 3. (H) Western blot analysis and (I) quantification of ALP, RUNX2, and COL1α1 in young/aged BMSCs co-cultured with different groups of BMDMs for 7d, n = 3. (J) Representative images of SA-β-gal staining of young/aged BMSCs co-cultured with different groups of BMDMs for 3d, scale bar, 50µm. (K) qPCR results of *IL-1b* and *IL-6* in young/aged BMSCs co-cultured with different groups of BMDMs for 3d, n = 3. Y-BMSCs, young BMSCs; A-BMSCs, aged BMSCs; shNC, BMDMs transfected with NC lentivirus; sh*Usp7*, BMDMs transfected with sh*Usp7* lentivirus; CON, control BMDMs; SEN, senescent BMDMs; Upper, upper well of transwell system; Lower, lower well of transwell system. Data are shown as the mean ± SD; ns, not significant, \**P* < 0.05, \*\**P* < 0.01, \*\*\**P* < 0.001, \*\*\*\**P* < 0.0001.

### USP7 inhibition enhanced aged BMSCs osteogenesis co-cultured with BMDMs

Next, we co-cultured USP7-depleted BMDMs with young/aged BMSCs. ALP/ARS staining indicated that USP7-depleted BMDMs significantly promoted osteogenesis in aged BMSCs (Fig. 2F). Moreover, USP7-depleted BMDMs significantly upregulated the mRNA and protein levels of osteogenic markers ALP, RUNX2, and COL1α1 in aged BMSCs (Fig. 2G-I). Regarding the senescence levels of aged BMSCs, SA-β-Gal staining indicated that USP7-depleted BMDMs significantly inhibited the senescence levels of aged BMSCs (Fig. 2J). Additionally, USP7-depleted BMDMs also inhibited the mRNA level of senescence-associated secretory phenotype (SASP) markers, *IL-1b* and *IL-6*, in aged BMSCs (Fig. 2K). Subsequently, we applied P5091 in the co-culture system. P5091 was found to improved osteogenesis by ALP/ARS staining (Fig. S4E). However, P5091 significantly inhibited COL1A1 expression in young BMSCs, while P5091 upregulated ALP, RUNX2, and COL1α1 in aged BMSCs (Fig. S4F). Regarding mRNA level, P5091 promoted osteogenic mRNA expression of *Alp, Runx2,* and *Collal* (Fig. S4G). The difference of mRNA and protein level of COL1α1 indicated that USP7 might played a post-transcriptional role in osteogenesis. Next, SA-β-Gal staining and qPCR also showed that P5091 also decreased the number of senescent cells and mRNA expression of *IL-1b*/*IL-6* in aged BMSCs co-cultured with BMDMs (Fig. S4H-I).

### USP7 inhibition promoted osteogenesis and senolysis of aged BMSCs

Next, we investigated the effects of USP7 depletion and P5091 on BMSCs without BMDMs. We successfully depleted USP7 expression in BMSCs (Fig. S5A-B). USP7-depleted aged BMSCs exhibited a trend of suppressed SASP pathways, and promoted osteogenic differentiation and apoptotic pathways (Fig. 3A-B). Furthermore, USP7 depletion inhibited cell viability of BMSCs significantly, while P5091 inhibited proliferation of aged BMSCs (Fig. 3C; Fig. S5C). Transwell migration assays further confirmed that USP7 depletion and P5091 promoted the migration of aged BMSCs (Fig. 3D; Fig. S5D). Regarding osteogenic capacity, ALP and ARS staining showed that USP7 depletion and P5091 enhanced osteogenesis of aged BMSCs, but P5091 inhibited osteogenesis of young BMSCs (Fig. 3E; Fig. S5E). Accordingly, USP7 depletion caused a significant reduction in the protein levels of ALP and COL1A1 in young BMSCs, while increasing these markers in aged BMSCs (Fig. 3F). P5091 significantly suppressed the expression of ALP and RUNX2 in young BMSCs, while substantially increasing the expression of osteogenic proteins in aged BMSCs (Fig. S5F). Additionally, USP7 depletion and P5091 significantly reduced the number of SA-β-gal-positive cells in aged BMSCs (Fig. 3G; Fig. S5G). USP7 depletion and P5091 also inhibited mRNA level and secretion level of IL-1β and IL-6 in aged BMSCs (Fig. 3H-I; Fig. S5H-I). Further, USP7 depletion and P5091 promoted cell death in aged BMSCs without affecting young BMSCs, which indicated that USP7 and P5091 could also function as senolytic drug in aged BMSCs (Fig. 3J-K; Fig. S5J-K).

**Fig. 3.**
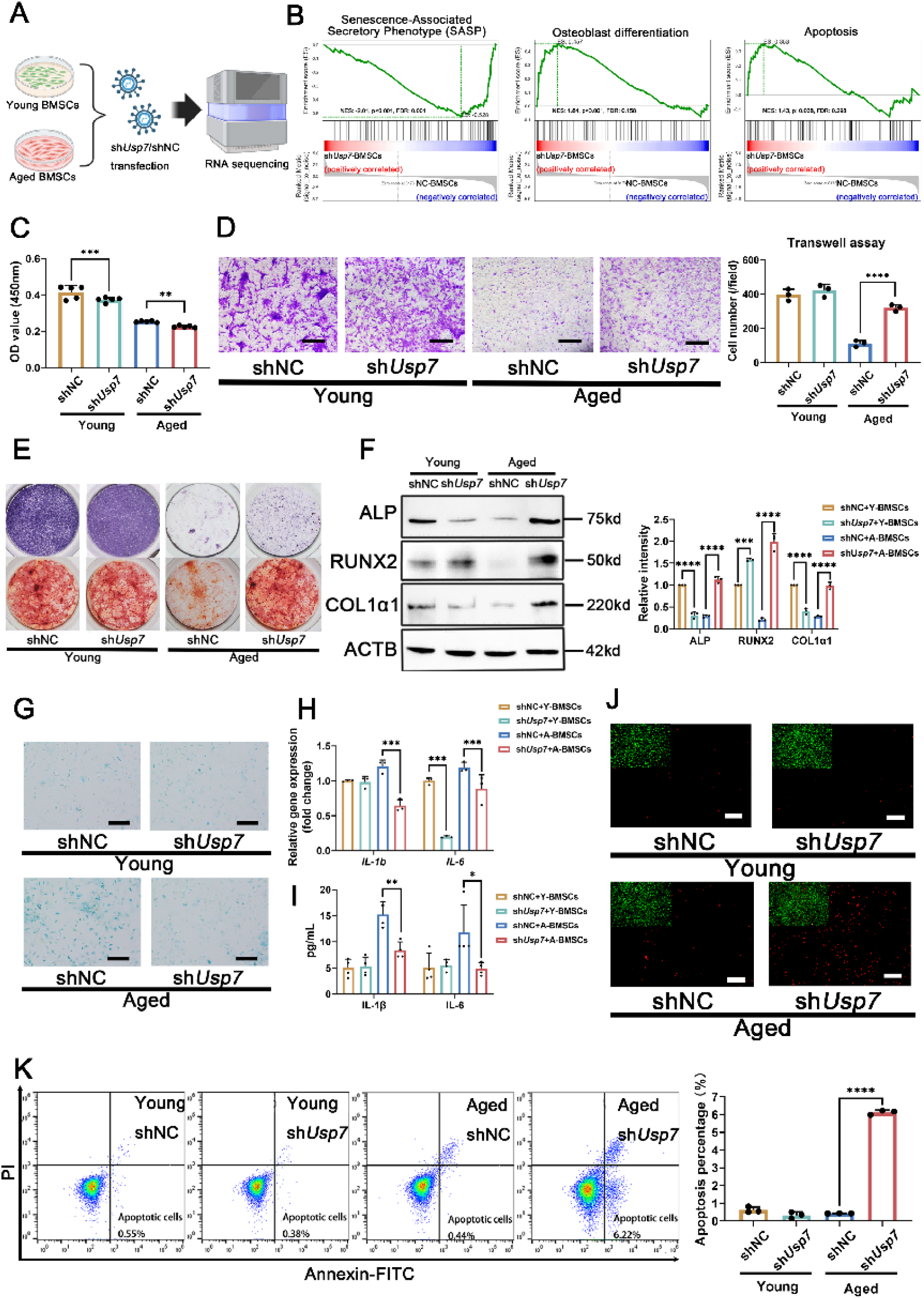
USP7 depletion promoted osteogenesis and senolysis in aged BMSCs. (A) schematic diagram showing mRNA-Seq analysis of young and aged BMSCs treated with shNC or sh*Usp7*. (B) GSEA showing decreased enrichment of SASP pathway-associated genes and elevated enrichment of osteoblast differentiation and apoptosis pathway-associated genes in USP7-depleted aged BMSCs. (C) CCK8 assay for determining cell viability of young/aged BMSCs transfected with shNC or sh*Usp7* lentivirus at 3d, n=5. (D) Representative images and quantitative analysis of migration capacity of young/aged BMSCs for 24 h, n = 3, scale bar, 50µm. (E) Representative images of ALP and ARS staining in different groups of BMSCs after initiating osteogenic induction for 7 and 14 d. (F) Western blot and quantification of ALP, RUNX2, and COL1α1 in different group of BMSCs after initiating osteogenic induction for 7 d, n = 3. (G) Representative images of SA-β-gal staining of different BMSCs, scale bar, 50µm. (H) qPCR results and (I) ELISA results of IL-1β and IL-6 in different groups of BMSCs, n = 3. (J) Representative images of live/dead staining of BMSCs after transfected with shNC or shUsp7 lentivirus for 3d, green color indicates live cells, red color indicates dead cells, scale bar, 150µm. (K) Flow cytometry and quantification for apoptosis in BMSCs after transfected with shNC or shUsp7 lentivirus for 3d. shNC, BMSCs transfected with shNC lentivirus; sh*Usp7*, BMSCs transfected with sh*Usp7* lentivirus. Data are shown as the mean ± SD; ns, not significant, \**P* < 0.05, \*\**P* < 0.01, \*\*\**P* < 0.001, \*\*\*\**P* < 0.0001.

### USP7 inhibition orchestrates EPSIN1/LRP1 pathway to enhance efferocytosis

Based on the mRNA-Seq results, KEGG pathway enrichment analysis revealed that the phagosome pathway was significantly upregulated in USP7-depleted senescent BMDMs (Fig. 4A). GSEA suggested that USP7 depletion upregulated efferocytosis capacity (apoptotic cell clearance) (Fig. 4B). The heatmap of core genes showed that LRP1 expression was significantly upregulated (Fig. 4C). In macrophages, EPSIN1 could inhibit efferocytosis by promoting the ubiquitination and degradation of LRP1 (Brophy et al. 2019). Through WB, we found that ubiquitination level was unchanged in different of groups of BMDMs. While USP7 depletion downregulated EPSIN1 and upregulated LRP1 protein levels in senescent BMDMs (Fig. 4D-E). Furthermore, after USP7 depletion, EPSIN1 protein levels gradually decreased over time, whereas overexpression of USP7 significantly delayed the degradation of EPSIN1 (Fig. S6A-B). Next, Co-IP experiments further validated the protein interaction between USP7 and EPSIN1 in BMDMs (Fig. 4F). Recombinant Flag-EPSIN1 and HA-USP7 proteins also demonstrated a similar interaction (Fig. S6C). Immunoprecipitating results showed that USP7 depletion led to an increased ubiquitination level of EPSIN1 (Fig. 4G). Simultaneously, overexpression of HA-USP7 reduced the ubiquitination level of EPSIN1 without affecting the overall ubiquitination level (Fig. S6D-E). This indicated that USP7 depletion might increase the ubiquitination and degradation of EPSIN, and LRP1 was spared from degradation, thereby promoting macrophage efferocytosis (Fig. S6F). Subsequently, we inhibited LRP1-mediated efferocytosis in BMDMs by using the LRP1 inhibitor RAP and si *Lrp1*. RAP inhibited the mRNA levels of efferocytosis-related genes such as *Cd36, Mertk*, and *Lrp1* which resulted in impeded efferocytosis (Fig. 4H-I; Fig. S7A). Additionally, USP7 depletion also enhanced the M2 polarization and inhibited pro-inflammatory cytokines of senescent BMDMs after efferocytosis, while RAP inhibited this effect (Fig. S7B-C). Further, USP7-depleted senescent BMDMs promoted osteogenic differentiation in aged BMSCs, while depleting LRP1 in BMDMs weakened this pro-osteogenic effect (Fig. S7D).

**Fig. 4.**
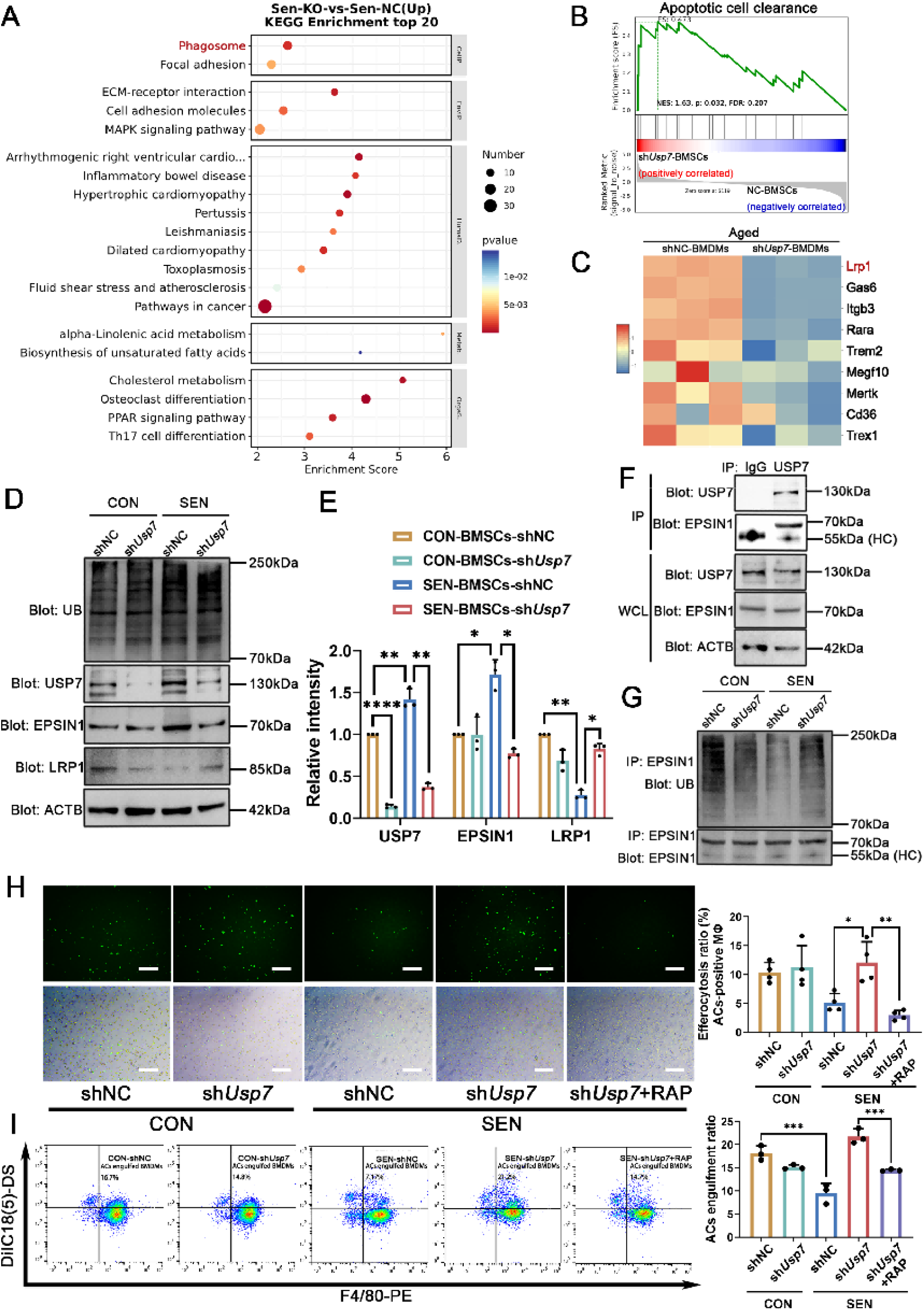
USP7 depletion promoted efferocytosis capacity in senescent BMDMs via EPSIN1/LRP1 pathway. (A) KEGG pathway enrichment analysis indicated an upregulated phagosome pathway in USP7-depleted senescent BMDMs. (B) GSEA showing elevated enrichment of apoptotic cell clearance pathway-associated genes. (C) Heatmap of representative apoptotic cell clearance-related genes in senescence and USP7-depleted senescent BMDMs. (D) Western blot and (E) quantification of ubiquitin, USP7, EPSIN1, and LRP1 in different group of BMDMs, n = 3. (F) Co-IP detecting the interaction between USP7 and EPSIN1. (G) Co-IP detecting the ubiquitous level of EPSIN1 in BMDMs transfected with shNC or sh*Usp7*. (H) Representative images and quantification of efferocytosis ratio of control/senescent BMDMs co-cultured with calcein-labeled apoptotic Jurkat cells for 24 h, n = 3, scale bar, 150µm. (I) Flow cytometry analysis for evaluating efferocytosis capacity of control/senescent BMDMs co-cultured with DiIC18(5)-DS-labeled apoptotic Jurkat cells for 24 h. DiIC18(5)-DS-positive and F4/80-postive cells indicated apoptotic cell-associated macrophages, n = 3. shNC, BMDMs transfected with NC lentivirus; sh*Usp7*, BMDMs transfected with sh*Usp7* lentivirus; CON, control BMDMs; SEN, senescent BMDMs; HC, heavy chain of IgG. Data are shown as the mean ± SD; ns, not significant, \**P* < 0.05, \*\**P* < 0.01, \*\*\**P* < 0.001, \*\*\*\**P* < 0.0001.

### USP7 inhibition promoted early stage of osseointegration in aged mice via efferocytosis

P5091 was used to inhibit USP7 in senile osteoporotic osseointegration in aged mice. P5091 treatment enhanced BIC and bone volume around implants in aged mice at 2-week and 4-week. The application of RAP attenuated the beneficial effect of P5091 (Fig. 5A-B). HE staining indicated that aged mice treated with P5091 showed bone structure formation around the implants at 2-week (Fig. S8A). Subsequently, we examined osteogenesis and osteoclastic activity around the implants using ALP and TRAP staining (Fig. 5C-D). ALP expression was increased in aged mice treated with P5091 compared to Veh and P50+RAP group at 2-week (Fig. 5E). Similarly, P5091 upregulated osteogenic genes in aged mice (Fig. S8B). Regarding osteoclastic activity, TRAP staining showed that at 2-week, P5091 downregulated the number of osteoclasts around the implants in aged mice. Additionally, the combined application of P5091 and RAP further reduced the number of osteoclasts. At 4 weeks post-implantation, the number of osteoclasts around the implants in young mice were significantly reduced, whereas aged mice still maintained a relatively high level (Fig. 5F). Regarding immune response, pro-inflammatory cytokines were upregulated in aged mice, and P5091 inhibited these in aged mice at 2-week and 4-week. However, RAP abrogated this effect (Fig. 5G). Besides, at 2 weeks post-implantation, in young mice, P5091 did not significantly alter the number of apoptotic cells, macrophage counts, or efferocytosis levels. However, in aged mice, P5091 significantly increased the number of apoptotic cells and macrophages, and promoted efferocytosis. While RAP did not alter the number of apoptotic cells, it significantly reduced the number of macrophages and suppressed efferocytosis (Fig. 5H-I). At 4 weeks post-implantation, both in young and aged mice, there were relatively fewer apoptotic cells and macrophages around the implants, and efferocytosis was also not evident (Fig. S9).

**Fig. 5.**
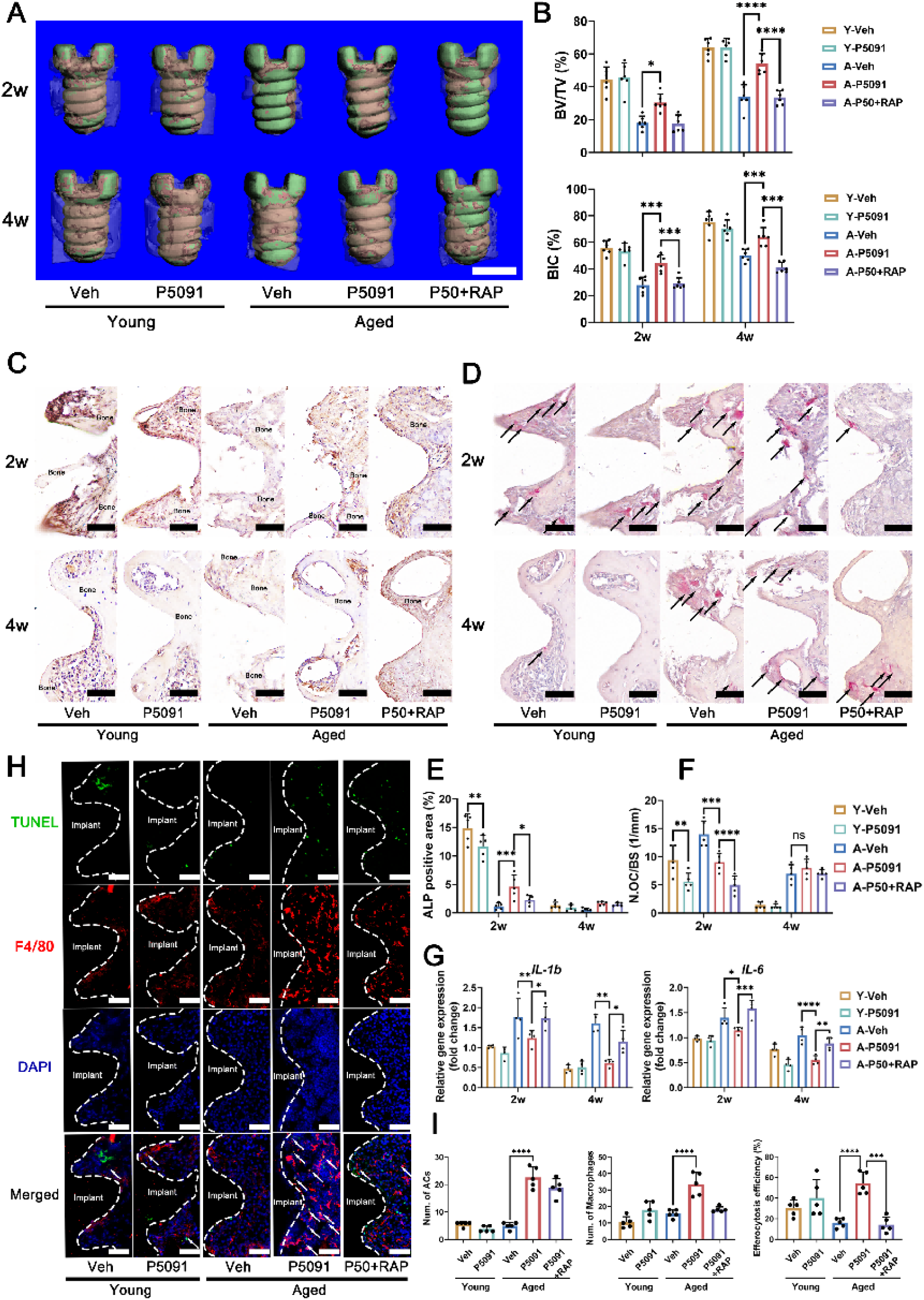
P5091 improved early stage of osseointegration in senile osteoporotic mice via efferocytosis. (A) Representative three-dimensional reconstruction images of alveolar bone around implants, scale bar. 600µm. (B) Quantitative MicroCT analysis of the 200µm region around the implants, n = 6. (C) Immunohistochemical staining of ALP around the implant, scale bar, 100µm. (D) TRAP staining around the implant, scale bar, 100µm. (E) Quantification of ALP-positive area in bone tissue around implants, n = 5. (F) Quantification of TRAP-positive osteoclasts in bone tissue around implants, n = 5. (G) qPCR results of *IL-1b* and *IL-6* expression in the bone tissue around implants, n = 4. (H) Representative immunofluorescence images of macrophages and apoptotic cells around the implants, and white arrows indicated the apoptotic cells engulfed by macrophages, scale bar, 100µm. (I) Quantification of apoptotic cell number, F4/80-positive macrophage number and efferocytosis efficiency (efferocytotic macrophages were determined as apoptotic cell-associated macrophages), n = 5. Veh, mice treated with vehicle (DMSO); P5091, mice treated with P5091; P50+RAP, mice treated with P5091 and RAP; Y, young mice; A, aged mice. Data are shown as the mean ± SD; ns, not significant, \**P* < 0.05, \*\**P* < 0.01, \*\*\**P* < 0.001, \*\*\*\**P* < 0.0001.

## DISCUSSION

In this study, we found that during the early stage of osseointegration, there was an abnormally high expression of USP7 around the implants in senile osteoporotic mice. However, Xie et al. argued that the femurs in OVX mice had a lower level of USP7 expression and USP7 lead to overactivated osteoclastogenesis (Xie et al. 2023). This discrepancy may stem from the use of different animal models. Although OVX mice exhibit an osteoporotic phenotype, they do not accurately replicate the senescent bone microenvironment. Lack of estrogen resulted in a higher bone turnover rate and osteoclastogenesis, however, senile osteoporosis is characterized by low bone turnover (Zhang et al. 2022). Therefore, we used senile osteoporotic male mice in this study and found that USP7 was overexpressed during osteoporotic osseointegration. Koyuncu et al. found that DUBs were abnormally upregulated in aged C. elegans, while inhibiting DUB prolonged their longevity (Koyuncu et al. 2021). Therefore, inhibiting USP7 may restore the normal cell function in age-related diseases.

Recent studies observed that diabetic wound tissues exhibited a higher level of USP7 expression and P5091 could improve wound healing via reducing SASP (Li et al. 2022; Zhang et al. 2024). Similarly, downregulating USP7/p300 pathway helps to inhibit senescence of multiple organs in chronic obstructive pulmonary disease (COPD) mice (Wang et al. 2023). However, Cui et al. argued that USP7 was vital for maintaining the normal life span via indirectly regulating DNA repair and oxidative stress (Cui et al. 2020). Therefore, it can be hypothesized that USP7 may have different functions in young and aged individuals.

Innate immune pathway and NLRP3 pathway were upregulated in the early stage of osseointegration (Chen et al. 2022). In this study, USP7 depletion and P5091 inhibited innate immune response and NLRP3 pathway in senescent BMDMs. Palazon et al. discovered that USP7 and USP47 directly maintained the NLRP3 pathway by preserving ASC speck formation (Palazón-Riquelme et al. 2018). Besides, USP7 also maintained NF-κB-mediated inflammatory pathway by stabilizing TRAF6 and p65 (Collins et al. 2015; Zhao et al. 2020). Therefore, USP7-depleted BMDMs may exhibit a weaker inflammatory response when exposed to aging stimuli. Li and colleagues discovered that in the bone marrow of aged mice, 80% of macrophages exhibit the M1 phenotype (whereas in young mice, M1 macrophages only make up 5%) (Li et al. 2021). Prolonged M1 polarization under inflammaging results in impaired BMSCs function by a robust amount of pro-inflammatory cytokines (IL-1β, IL-6, MMP2) (Lee et al. 2020). In this study, USP7-depleted senescent BMDMs showed a reduced level of inflammatory cytokines, which alleviated their inhibitory effect on osteogenesis. As a result, USP7 inhibition might inhibit inflammaging of senescent BMDMs, thereby improve osteogenesis of aged BMSCs.

On the other side, aged BMSCs also contributed to inflammaging and M1 polarization by secreting SASP factors such as IL-1β and IL-6 (Lee et al. 2020). Therefore, we investigated how USP7 inhibition affected aged BMSCs. We observed that USP7 depletion, inhibited cell viability of BMSCs while increased osteogenic capacity in aged BMSCs. Similarly, Kim et al. demonstrated that after USP7 depletion, the ubiquitination levels of NANOG and SOX2 increased, subsequently inhibiting self-renewal capacity and multipotency of BMSCs (Kim et al. 2022). Consequently, USP7 plays a different role between young and aged BMSCs. In this study, we discovered that USP7 depletion promoted apoptosis and osteoblast differentiation in aged BMSCs, while SASP pathway was inhibited. He et al. found that USP7 inhibition could enhance p53/p21 pathway via degradation of MDM2, which lead to the senolysis of senescent cells (He et al. 2020). Therefore, senolytic therapies by USP7 inhibition in aged BMSCs may also promote osteogenesis and osseointegration in aged individuals.

While senolysis-mediated apoptosis of senescent cells aids in tissue repair, the efficient clearance of apoptotic cells is equally important for successful wound healing (Mehrotra et al. 2022). Phagocytosis of apoptotic cells is a crucial process for macrophages to transition from the M1 phase to the M2 phase during the wound proliferation stage, which is also known as efferocytosis (Gerlach et al. 2021). In aging individuals, macrophages may experience impaired efferocytosis functions because of impaired MerTK function (Rymut et al. 2020). This may lead to ineffective clearance of apoptotic cells and prolonged inflammation stage in early osseointegration. In this study, we observed that USP7 depletion improved the impaired efferocytosis via upregulation of LRP1 in senescent BMDMs. LRP1 is a key receptor involved in mediating efferocytosis, which works with AXL and CD36 to recognize “Eat me” signals (Doddapattar et al. 2022). LRP1 was found to play a vital role in osteoclast-osteoblast interactions (Lu et al. 2018). Vi et al. found that macrophage LRP1 was crucial for fracture repair but did not propose related signaling mechanisms (Vi et al. 2018). Therefore, LRP1-mediated efferocytosis may play a crucial role in osteoimmunology, although corresponding studies are lacking.

In the *in vivo* experiments, since knockout of USP7 might lead to embryonic lethality, and tamoxifen required for CreERT2 can affect bone metabolism, we therefore used P5091 to inhibit USP7 in mice (Chen et al. 2023; Kon et al. 2010). We observed that P5091 induced senolysis and osseointegration at 2-week in aged mice. This suggested that the senolysis induced by P5091 contributes to osseointegration in aged mice. Farr et al. utilized dasatinib and quercetin, a classic senolytic therapy, to eliminate senescent cells in aged mice and alleviate of osteoporosis (Farr et al. 2017). Besides, P5091 improved osseointegration via enhancing efferocytosis. We found that efferocytosis was impeded in aged mice at 2-week while P5091 upregulated macrophage efferocytosis. However, the apoptotic cell-associated macrophages could be merely observed at 4-week. This may be because a significant amount of cell apoptosis and immune clearance occurs primarily during the early stages of healing. We also observed that RAP inhibited efferocytosis promoted by P5091, suggesting that LRP1 played a vital role in macrophage efferocytosis of aged mice. Simultaneously, we found that P5091 inhibited pro-inflammatory cytokines, while inhibiting efferocytosis via RAP upregulated these in aged mice. However, RAP significantly reduced osteoclast number at 2-week. Kohara et al. found that LRP1 deficiency leads to inhibited osteoclastogenesis (Kohara et al. 2020). And LRP1 deficiency also impedes osteoblast function and osteoblast-osteoclast interaction by disrupting PDGF-RANKL pathway (Bartelt et al. 2018). Consequently, the effect of RAP on inhibiting osteoclast function during osseointegration in aged mice may be partially attributed to its direct action on osteoclasts. Therefore, therapies targeting efferocytosis may improve senile osteoporotic osseointegration, which was related to reduced immune response mediated by efferocytosis.

In conclusion, we first identified an abnormally high expression of USP7 in the peri-implant tissues and aged BMSCs/BMDMs. USP7 inhibition could alleviate inflammaging in senescent BMDMs and induce apoptosis in aged BMSCs. Interestingly, we discovered that USP7 inhibition promotes the efferocytosis of senescent BMDMs via EPSIN1/LRP1 pathway, thereby enabling efficient clearance of senescent cells. P5091, a selective inhibitor of USP7, could enhanced early osseointegration in aged mice via efferocytosis and senolysis.This study underscores that USP7 may be a potential target for promoting early osseointegration in elderly patients with osteoporosis.

## Declaration of Conflicting Interests

The authors declared no potential conflicts of interest with respect to the research, authorship, and/or publication of this article.

## Acknowledgement

We acknowledge the funding from the National Natural Science Foundation of China (81970966, 82001084), Natural Science Foundation of Sichuan Province (2023NSFSC0574, 2023NSFSC1513).

## Author Contributions

F. Zhou, contributed to conception and design, data acquisition, analysis, and interpretation, drafted and critically revised the manuscript; Z. Wang, H Li, D. Wang, contributed to conception and design, data interpretation, critically revised the manuscript; Z. Wu, B. F, contributed to conception, data analysis, drafted and critically revised the manuscript; Q. Wang, W. Luo, G. Zhang, contributed to conception, data acquisition, drafted and critically revised the manuscript; Y. Xiong, Y. Wu, contributed to design, data acquisition and interpretation, drafted and critically revised the manuscript. All authors gave their final approval and agree to be accountable for all aspects of the work.

## Data and materials availability

All data are available in the main text or the supplementary materials upon reasonable requests

## Notes

### Competing Interest Statement

The authors have declared no competing interest.

### Summary of Updates

Add figures and revise the manuscript

