## Supplementary material for "USP7 Inhibition Promotes Early Osseointegration in Senile Osteoporotic Mice": Methods and Materials

**Affiliations：**

***Correspondence should be addressed to:**

Yingying Wu, D.D.S, Ph.D.

**Appendix Materials and Methods**

**In vitro experiments**

**Cell culture and co-culture of BMSCs-BMDMs**

Young and aged BMSCs were extracted from femurs and tibia in 6-month and 18-month mice, which were cultured in the growth medium consisting of low-glucose DMEM (Gibco), supplemented with 10% FBS (Gibco), and 1% penicillin and streptomycin. BMDMs were extracted from 6-month mice, and plated in pre-cooled prepared bone marrow macrophage differentiation medium containing high-glucose DMEM (Gibco), 10% FBS (Gibco), 1% penicillin/streptomycin, and 20 ng/mL monocyte-colony stimulating factor (M-CSF, PeproTech). When inducing senescent BMDMs, conditional media (CM) from aged BMSCs was used to culture BMDMs. CM from aged BMSCs was mixed with macrophage differentiation medium in a 1:3 ratio. To inhibit USP7 and LRP1 in cells, P5091 (MCE) was dissolved in DMSO and applied in BMSCs and BMDMs at the concentration of 5μM, while receptor-associated protein (RAP, Abcam) was dissolved in PBS and applied in BMDMs at the concentration of 150nM.

As for the co-culture of BMSCs-BMDMs, in the indirect co-culture system, 0.4 μm 24-well Transwell inserts were utilized to co-culture BMSCs and BMDMs. For the direct co-culture system, a 24-well plate was used to co-culture BMSCs and BMDMs. A total of 60,000 BMSCs and 60,000 BMDMs were simultaneously seeded into the 24-well plate, and 500 μL of growth medium containing 20 ng/mL M-CSF was added.

**Transfection of BMSCs and BMDMs**

BMDMs and BMSCs were transfected with a lentiviral psi-LVRU6P vector expressing Usp7 shRNA or control shNC (shNegativeControl, GeneCopoeia). When BMSCs or BMDMs reached approximately 70-90% confluence, they were subjected to transfection at multiplicity of infection (MOI) of 50 with polybrene (5 mg/mL, Sigma). The transfection process took place at 37°C in a 5% CO2 environment for 24 hours. Subsequently, the supernatant was removed, and normal growth medium was added. After 2 days, puromycin (Sigma) was used to select shRNA-induced cells. As for siRNA transfection, si*Lrp1* (GeneCopoeia) was transfected to BMDMs with application of EndoFectin Max (GeneCopoeia). After 18h, BMDMs were rinsed by PBS and cultured with complete medium for 48h. Then BMDMs transfected with si*Lrp1* were used for subsequent experiments.

**mRNA-seq**

The transcriptome sequencing and subsequent analysis were conducted by OE Biotech Co., Ltd. (Shanghai, China). Briefly, total RNAs were extracted utilizing the TRizol reagent (Invitrogen). The purity and concentration of the RNA were determined with a NanoDrop 2000 spectrophotometer (Thermo Scientific). Subsequently, libraries were prepared in accordance with the manufacturer's protocol using the VAHTS Universal V6 RNA-seq Library Prep Kit. The libraries were subjected to sequencing on an Illumina Novaseq 6000 platform, yielding 150 bp paired-end reads. Quality control of the mRNA-seq data was carried out using FastQC (v0.11.5) and the FASTX toolkit (0.0.13). Alignment to the Mus musculus reference genomes was performed using HISAT2 (v.2.0.4). Differentially expressed genes between shUsp7 and shNC groups were identified using the DEseq2 package (version 1.34.0) if they met the criteria of Q value < 0.05 and a fold change greater than 2 or less than 0.5.

**Cell proliferation assay**

The cell proliferation of BMSCs and BMDMs was evaluated using Cell Counting Kit-8 (CCK-8, Beyotime) assay. A mixture containing 100 μL of 10% CCK-8 assay reagent was added to each well. Following incubation in the absence of light for 1.5 hours, the optical density (OD) was determined at 450 nm using a Varioskan Flash microplate reader (Thermo Scientific).

**Migration assay**

BMSCs were initially plated at a density of 2 × 10^4^ cells per well in 8 μm 24-transwell plates (Corning). Following a 24-hour incubation period, BMSCs present in the transwell chambers were fixed for 30 minutes by 4% paraformaldehyde (Biosharp), stained with 0.1% w/v crystal violet (Beyotime) for an additional 45 minutes, and subsequently visualized using an inverted microscope (Leica).

**RNA extraction and quantificational real-time PCR (qRT-PCR)**

Total RNA was extracted from the alveolar bone tissue around implants and treated BMSCs/BMDMs following the manufacturer's guidelines, employing a total RNA isolation kit from Vazyme. Bone tissue around implants was first separated from maxilla and immersed in liquid nitrogen, which were were ground into a paste using a mortar and pestle. Subsequently, RNA samples underwent reverse transcription into cDNA utilizing the HiScript® III All-in-one RT SuperMix, ideally suited for qPCR (Vazyme). Quantitative real-time polymerase chain reaction (qRT-PCR) was carried out using Taq Pro Universal SYBR qPCR Master Mix (Vazyme). A list of the primers employed is provided in Supplementary Table 1. The relative gene expression levels were determined using the 2^-△△^CT method, normalizing to β-actin as the internal reference gene.

**Western blot**

The cell protein of treated BMDMs and BMSCs was extracted using RIPA buffer (Pierce) while maintained on ice. As for alveolar bone tissue around implants, the alveolar bone around the implant was first separated and then immersed in liquid nitrogen. The bone was then placed into a centrifuge tube containing grinding beads and ground to obtain a protein lysate. Following this, the samples were denatured at 95 ℃ for 10 minutes in SDS loading buffer and subsequently separated on 10% SDS–polyacrylamide gels (Beyotime). The separated proteins were then transferred onto polyvinylidene difluoride (PVDF, Millipore) membranes activated with methanol. After 5% BSA was applied to the membranes and allowed to incubate for 2 hours, the membranes were exposed to primary antibodies (listed in Supplementary Table 2) overnight at 4 ℃. The following day, the membranes were subjected to incubation with corresponding horseradish peroxidase (HRP)-conjugated secondary antibodies, followed by an additional incubation for 1–2 hours at 37 ℃. The detection of antibody-antigen complexes was achieved using gel imaging systems (Bio-Rad). The quantification of band densities was performed using ImageJ (National Institutes of Health) and subsequently normalized to the control.

**Osteogenic induction of BMSCs and ALP/ARS staining**

When BMSCs reached 70% confluency, culture medium was replaced with mineralization medium [growth medium supplemented with ascorbic acid (50μg/mL), b-glycerophosphate disodium (10 mM), and dexamethasone (100nM)].

As for ALP staining, after 7 days of osteogenic induction, cells were first fixed by 4% paraformaldehyde (Biosharp) for 15 min, and then stained with a BCIP/NBT Alkaline Phosphatase Color Development Kit (Beyotime) for 24 h.

As for ARS staining, after 14 days of osteogenic induction, cells were first fixed by 4% paraformaldehyde (Biosharp) for 15 min, and then stained with Alizarin Red S solution (pH 4.2, Sigma) for 5 min.

**Senescence-associated β-galactose staining**

Cellular senescence was evaluated using senescence-associated β-galactosidase (SA-β-gal) staining kits (Beyotime). BMSCs were rinsed twice with PBS and subsequently fixed with a fixative solution for SA-β-gal staining for 15 minutes at room temperature. Then cells were washed three times with PBS and then incubated with the SA-β-gal staining solution at 37 °C for 24 h. The following day, cells were examined under an optical microscope.

**Enzyme-Linked immunosorbent assays (ELISA)**

The cell culture supernatant of BMDMs and BMSCs was harvested following the provided guidelines. The levels of IL-1β, TNF-α, and IL-6 in the supernatant were quantified utilizing ELISA kits (Elabscience).

**Flow cytometry**

To characterize BMDMs，BMDMs were prepared in the staining buffer (BD Biosciences)**.** To prevent nonspecific binding, BMDMs were pre-incubated with FC block (BD Biosciences, 1:200) for 20 minutes. They were then incubated with the following antibodies on ice for 30 minutes: PE Rat Anti-Mouse F4/80 (BD Biosciences, 1:100), FITC Rat Anti-CD11b (BD Biosciences, 1:100). After incubation, the cells underwent three washes with staining buffer and were subsequently resuspended in staining buffer. After the last wash and centrifugation, the cells were once again resuspended in staining buffer for analysis using Attune Nxt Flow Cytometer (Thermo Scientific).

To characterize BMSCs, BMSCs were suspended in staining buffer and stained respectively with BV605 Rat Anti-Mouse CD45 (BD Biosciences, 1:200), FITC Mouse Anti-Rat CD90/Mouse CD90.1 (BD Biosciences, 1:200), PE Rat Anti-Mouse CD73 (BD Biosciences, 1:100), BV421 Rat Anti-Mouse CD44 (BD Biosciences, 1:100), FITC Hamster Anti-Rat CD29 (BD Biosciences, 1:100), andAlexa Fluor™ 488 Rat Anti-Mouse CD34 (BD Biosciences, 1:200). After the last wash and centrifugation, the cells were once again resuspended in staining buffer for analysis using Attune Nxt Flow Cytometer (Thermo Scientific).

To detect apoptosis level, BMSCs treated with P5091 for 3 days and *shUsp7* were prepared in staining buffer and stained with 1 μL FITC Annexin V and PI (APEBIO) for 30 minutes. After incubation, the cells were washed by PBS for 3 times. After the last wash and centrifugation, the cells were once again resuspended in staining buffer for analysis using Attune Nxt Flow Cytometer (Thermo Scientific).

As for efferocytosis and efferocytosis-induced M2 polarization, Jurkat cells (SCC-121815, Solarbio) were first labeled by DiIC18(5)-DS (MCE), then exposed to UV light (150 mJ/cm2) for 15 minutes with an open lid. Then apoptotic Jurkat cell were incubated with BMDMs for 24 h. After that, BMDMs were suspended in the staining buffer (BD Biosciences). To prevent nonspecific binding, BMDMs were pre-incubated with FC block (BD Biosciences, 1:200) for 20 minutes. They were then incubated with either of the following antibodies on ice for 30 minutes: PE Rat Anti-Mouse F4/80 (BD Biosciences, 1:100) Alexa Fluor® 647 Rat Anti-Mouse CD206 (BD Biosciences, 1:50). After incubation, the cells underwent three washes with staining buffer and were subsequently resuspended in staining buffer. After the last wash and centrifugation, the cells were once again resuspended in staining buffer for analysis using Attune Nxt Flow Cytometer (Thermo Scientific). FACS data analysis was conducted using FLOWJO™ Software.

**Live/dead staining**

Calcein-AM (Beyotime) and 7-Aminoactinomycin D (7-AAD, Beyotime) were employed for concurrent fluorescence assessment. First, Calcein-AM (5 μM) was added into the culture medium for 30 min. Then, after cells were washed by PBS for 3 times, 7-AAD (5 μM) were added into the culture medium for 10 min. The distinction between living (green) and deceased (red) cells was ascertained using a fluorescence microscope and Image J software (National Institutes of Health) for cell quantification.

**Efferocytosis assay (Immunofluorescence staining)**

First, Jurkat cells (SCC-121815, Solarbio) were labeled with 5 µM Calcein-AM (Beyotime). Following a 2-hour incubation, Cells were washed for 3 times and subjected to UV irradiation (150 mJ/cm2) with an open lid. Subsequently, they underwent an additional 2-hour incubation before being introduced in a 1:1 ratio to BMDMs for 1 h. After 3 times of gentle washing, BMDMs were considered positive for phagocytosed green apoptotic Jurkat cells if they contained clusters of green dots measuring >3 µm in size. Quantification was performed based on 3 images by Image J software (National Institutes of Health).

**Co-IP**

Treated BMDMs were lysed using Pierce IP Lysis Buffer (Thermo Fisher Scientific) following the provided instructions. Antibodies against USP7 (1:50, 31687, Cell Signaling Technology), EPSIN1 (1:100, ab86064, Abcam), HA (1:100, ab9110, Abcam; 1:100, M20013, Abmart), and Flag (1:100, ab205606, Abcam; 1:100, M20008, Abmart) were added to the lysates overnight at 4℃, and protein A/G beads were added into the mixture for another 4 h. After centrifugation, the pellet was collected and washed three times with lysis buffer. Subsequently, it was resuspended in RIPA buffer and boiled with SDS loading buffer at 95 ℃ for 10 minutes. The boiled samples were centrifuged again, and the supernatant was utilized for subsequent Western blot.

**In vivo experiments**

**Mice**

All animal experiments were approved by Subcommittee on Research and Animal Care (SRAC) of Sichuan University (WCHSIRB-D-2020-163). Young (3-month-old) and aged (18-month-old) male wild-type C57BL/6J mice were purchased from Chengdu Dossy Experimental Animals Co. Ltd. The mice were housed in SPF facilities with a 12 h light/dark cycle and provided with sufficient food and water. A total of 120 mice were used in this study. The sample size in different experiments was based on former studies and pre-experiments which was analyzed by One-way ANOVA F test. We randomized mice by body weight using the random function in EXCEl. The mice were randomly divided into 5 groups under double blindness: young mice treated with vehicle, young mice treated with P5091, aged mice treated with vehicle, aged mice treated with P5091, and aged mice treated with P5091 and RAP. Mice without systemic diseases and peri-implant inflammation in each group were included in the subsequent experiments and analyses. During experiment periods, no mice were excluded. USP7 inhibitor P5091 was dissolved in PBS containing 4% DMSO. Mice were injected daily i.p. with injections of vehicle (4% DMSO in PBS) or P5091 (5 mg/kg) or receptor-associated-protein (RAP, 20mg/kg) for 4 weeks based on former studies. All animal experiments described above were conducted in accordance with the ARRIVE guidelines.

**Implant surgery**

All mice were were anesthetized via intraperitoneal injection of ketamine at 80 mg/kg (Sigma Chemical) and xylazine at 10 mg/kg (Sigma Chemical). Following oral saline cleaning, a 2 mm incision was made anterior to the left first upper molars to mark the implant site. The implant site was prepared under saline cooling using a surgical motor (NSK-Nakanishi). Subsequently, the pilot hole was adjusted to accommodate the titanium implant (WEGO) using a drill. The titanium implant was then screwed into the implant bed with a screwdriver. Following the surgery, the mice were provided with a soft diet and were injected daily i.p. with injections of vehicle (4% DMSO in PBS) or P5091 (5 mg/kg) for 4 weeks.

**MicroCT**

The harvested femur and maxillary bones were first fixed in 4% paraformaldehyde for 24 h. Micro-CT analysis was carried out using the Hiscan XM Micro-CT system (Hiscan Information Technology Co., Ltd.), with a spatial resolution of 10 μm and utilized the following settings: 80 kV, 100 μA, and 50 ms integration time per step. As for femur, the region of interest (ROI) was defined as the region located 700 μm below the growth plate of the femur. Bone morphological parameters, including, bone volume per total volume (BV/TV), trabecular number (Tb.N), and trabecular thickness (Tb.Th), cortical bone thickness (Ct.Th), and cortical bone area (Ct.Ar) were assessed within the ROI. As for the maxilla, the region of interest, composed of a radius of 200 μm around the implants, was selected for the reconstruction of alveolar bone, encompassing the central 50% portion of the implants. Parameters including BV/TV and bone-implant contact (BIC) were evaluated within this defined region.

**Histomorphometric/immunofluorescence staining analysis**

For histological analyses, the maxillary bones were initially fixed in 4% paraformaldehyde at 4 °C for 24h and subsequently transferred to PBS at 4 °C.

As for immunofluorescence staining and immunohistochemical staining, the maxillary bones were decalcification in EDTA (pH 7.2, Biosharp) for 6 weeks, allowing the bone to become pliable. The decalcified tissue were then embedded in paraffin, and 4-μm sections were obtained for subsequent staining.

As for immunofluorescence staining, the following primary antibodies and fluorescent dyes were utilized in this study: anti-F4/80 (Abcam, ab6640, 1:100), DAPI (Beyotime), and TUNEL (Beyotime). The detection of the primary antibodies was carried out using the corresponding Alexa Fluor-conjugated secondary antibodies (Abcam). All tissue sections were examined under a fluorescent microscope (Leica).

As for immunohistochemical staining, the sections were stained by ALP antibody (ET1609-68, Huabio) and USP7 antibody (A300-033A, ThermoFisher). The positive-expression area of ALP wa calculated by IHC plugins in imageJ (National Institutes of Health). The ratio of ALP-positive area was calculated by the pixels of positive-expression area of ALP and the pixels of whole bone tissue in images. TRAP staining was performed according to the instructions provided in the Beyotime TRAP Staining Kit (P0332). The number of osteoclasts was calculated by imageJ (National Institutes of Health).

**Statistical analysis**

Statistical analyses were conducted using GraphPad Prism version 8.0 and SPSS 22.0 software. The data are expressed as means with standard deviation (SD). The normality of data was first verified by Shapiro–Wilk test. Two-tail student's t-test was employed to compare two groups, while one-way analysis of variance (ANOVA) was applied when analyzing three or more than three groups. For multiple comparisons, the Bonferroni's multiple comparisons test was utilized. Sample sizes are specified in the figure legends. Statistical significance was defined as **P*< 0.05, ***P* < 0.01, ****P* < 0.001, and *****P* < 0.0001.

**Appendix Results**

**18-month-old male mice showed an osteoporotic phenotype in the femur and alveolar bone**

In the femurs of male mice at 6, 12, and 18 months, MicroCT results showed that the 18-month-old mice exhibited a significant osteoporotic phenotype (Fig. S1A). Compared to the 6-month-old mice, the 18-month-old mice had significantly reduced bone volume and trabecular bone thickness with increased trabecular bone separation in the cancellous bone (Fig. S1B). Additionally, the cortical bone thickness and area in the 18-month-old mice were also significantly decreased (Fig. S1C). Similarly, the interradicular bone around the first maxillary molar showed reduced bone volume, trabecular bone thickness, and higher trabecular bone separation in 18-month-old mice, suggesting that 18-month-old mice also showed an obvious osteoporotic phenotype in alveolar bone (Fig. S1D-E).

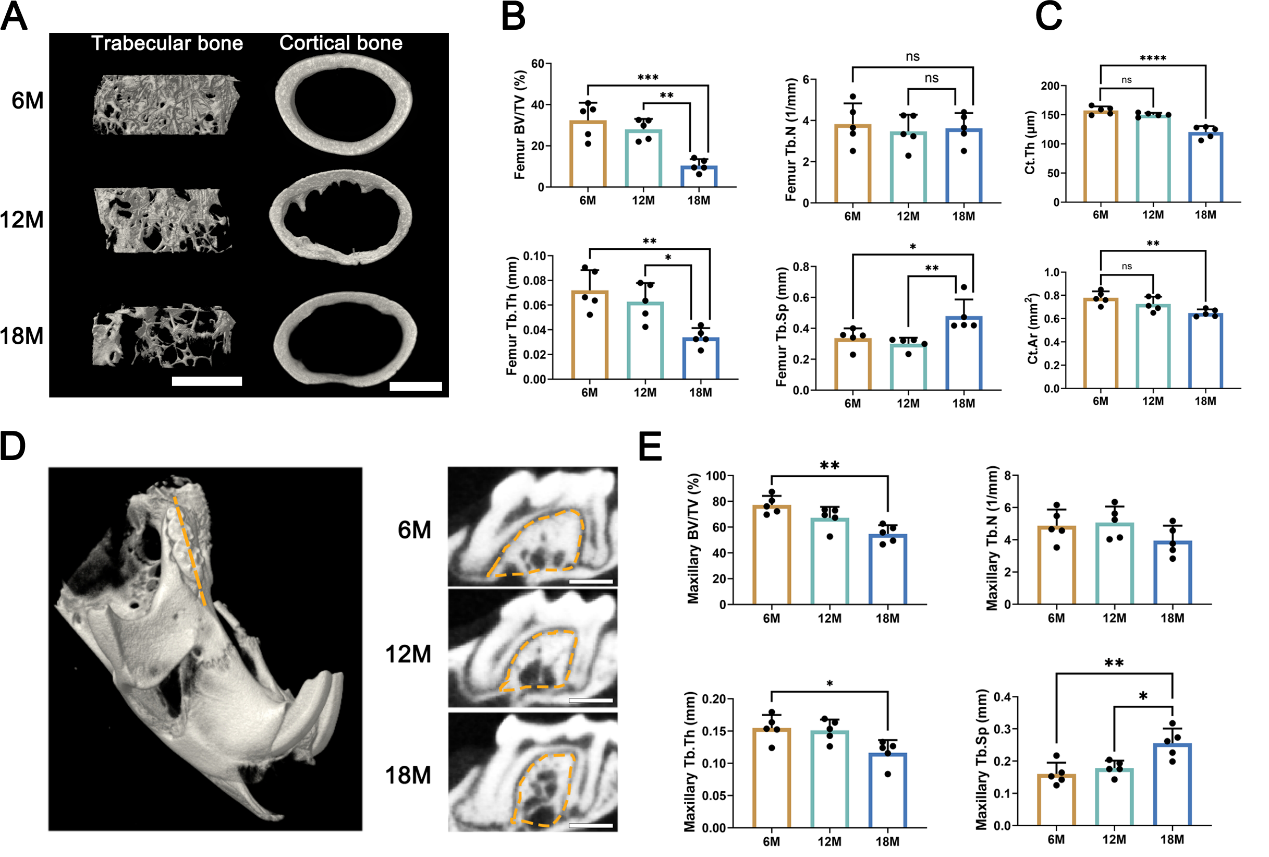

Appendix Fig. 1 18-month-old mice exhibited an obvious osteoporotic phenotype in both the femur and alveolar bone. Different age of male mice (6-month, 12-month and 18-month) were employed. (A) Three-dimensional reconstruction image of cancellous and cortical bone in femurs at the position 700μm below the growth plate, scale bar, 500μm. (B) Quantitative MicroCT analysis of cancellous bone, n = 5. (C) Quantitative MicroCT analysis of cortical bone, n = 5. (D) Three-D dimensional reconstruction of maxillary alveolar bone in mice. The orange dashed lines indicated the orientation of sagittal micro–computed tomographic sections of interradicular bone around the first molar. (E) Quantitative MicroCT analysis of interradicular bone around the first molar. n = 5. BV/TV, bone volume per tissue volume; Tb.N, trabecular number; Tb.Th, trabecular thickness; Tb.Sp, trabecular separation; Ct.Th, cortical bone thickness; Ct.Ar, cortical bone area; BIC, bone-implant intact. Data are shown as the mean ± SD; ns, not significant, **P* < 0.05, ***P* < 0.01, ****P* < 0.001, *****P* < 0.0001.

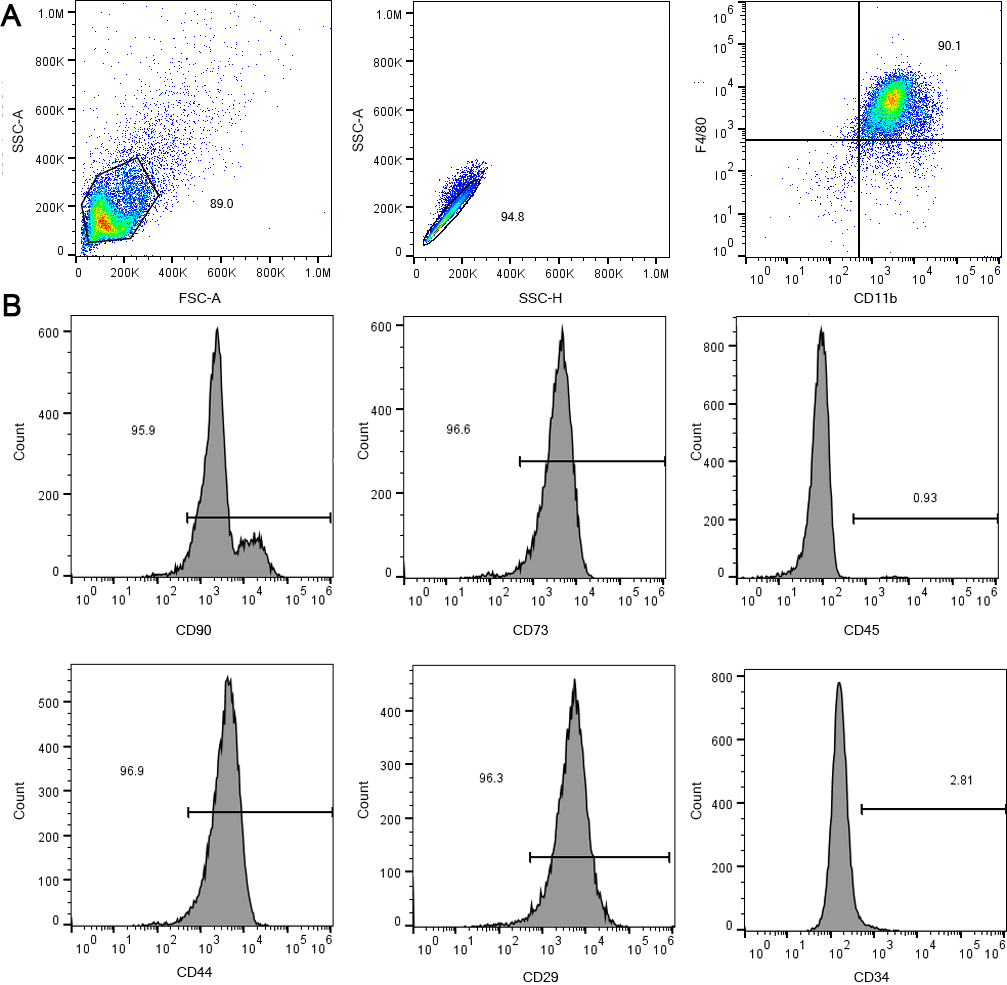

**Appendix Figure 2.** Characterization of BMDMs and BMSCs. (A) Flow cytometry showed that over 90% of the cells were CD11b/F480 positive which were the makers for murine macrophages. (B) Flow cytometry showed that over 95% of the cells were positive for CD90, CD73, CD44, and CD29 while negative for CD45 and CD34, which exhibited the characteristics of murine BMSCs.

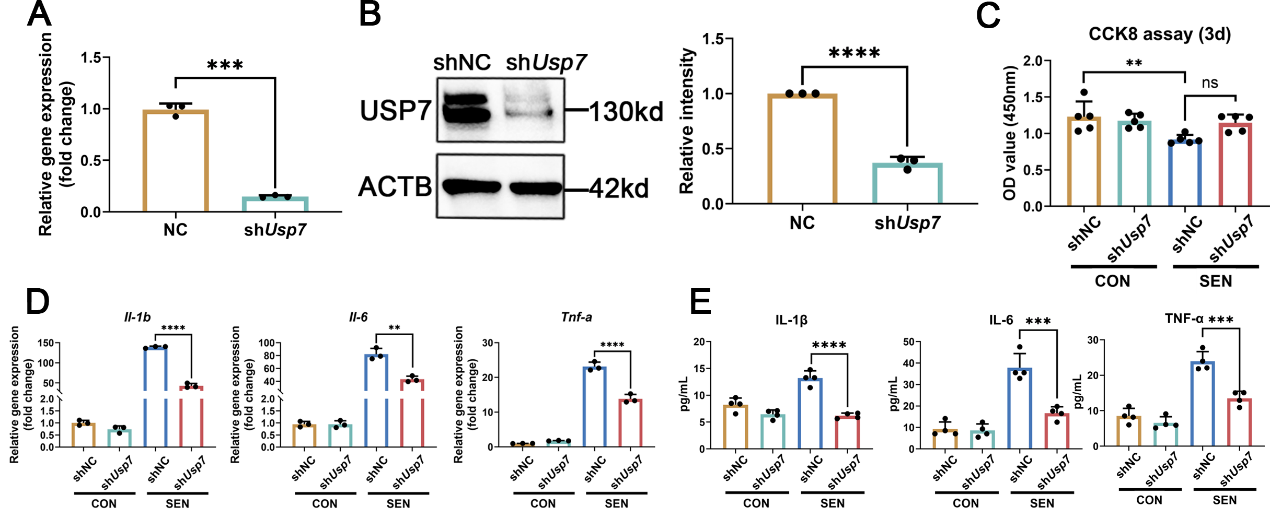

**Appendix Figure 3.** Generation of USP7-depleted BMDMs and USP7 depletion inhibited innate immune response. (A) qPCR showing downregulated mRNA level of USP7 in BMDMs transfected with sh*Usp7*, n = 3. (B) WB analysis showing downregulated protein level of USP7 in BMDMs transfected with sh*Usp7*, n = 3. (C) CCK8 assay determining cell viability of control/senescent BMDMs at 3d after transfected with sh*Usp7*, n = 5. (D) qPCR results of *IL-1b, IL-6*, and *Tnf-a* mRNA expression in different groups of BMDMs, n = 3. (E) ELISA results of IL-1β, IL-6, and TNF-α secretion level in different groups of BMDMs, n = 4. shNC, BMDMs transfected with NC lentivirus; shUsp7, BMDMs transfected with shUsp7 lentivirus; CON, control BMDMs; SEN, senescent BMDMs. Data are shown as the mean ± SD; ns, not significant, **P* < 0.05, ***P* < 0.01, ****P* < 0.001, *****P* < 0.0001.

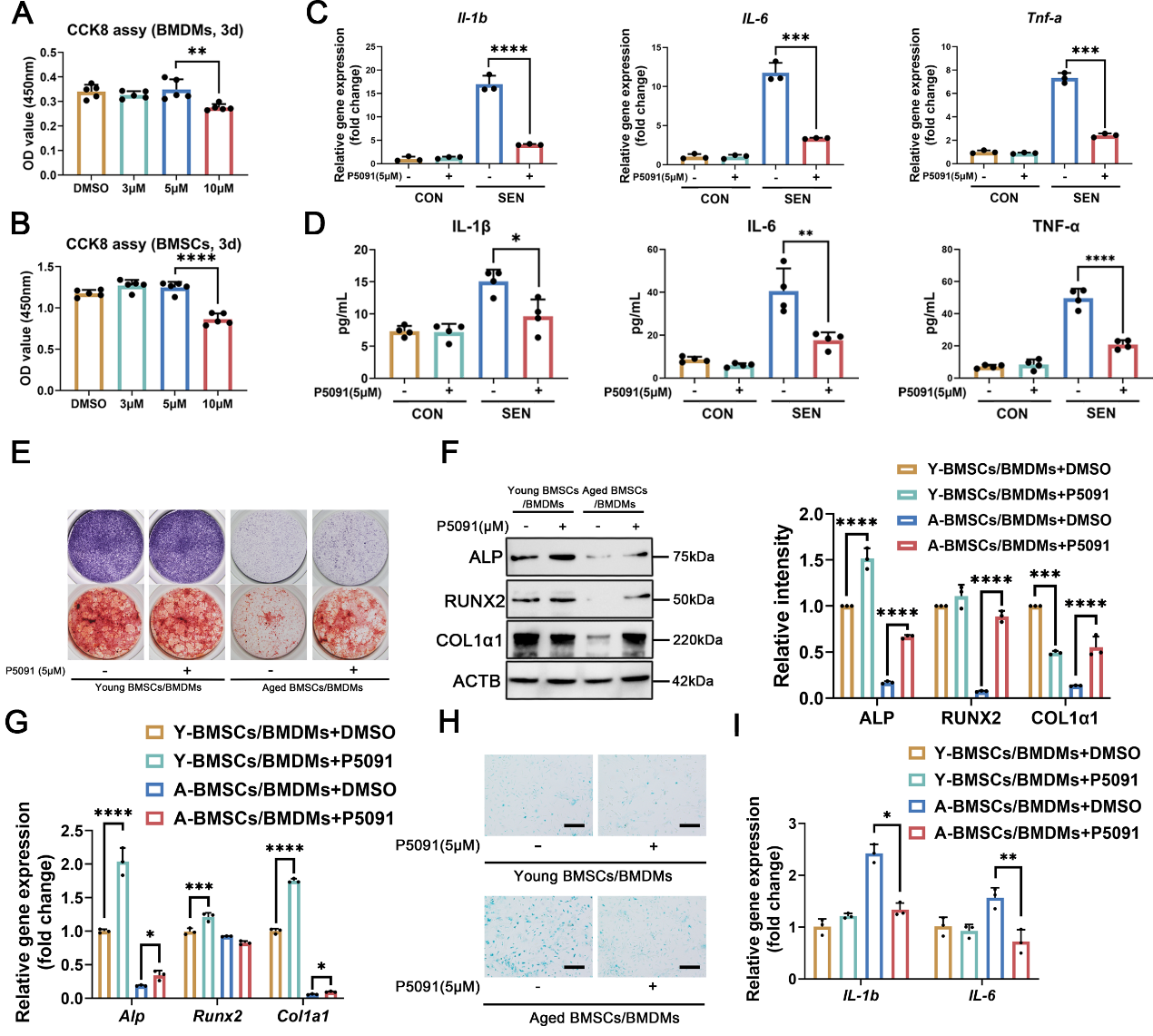

**Appendix Figure 4.** P5091 inhibited inflammation in senescent BMDMs and promoted osteogenesis of aged BMSCs co-cultured with BMDMs. CCK8 demonstrating the changes in cell viability of BMDMs (A) and BMSCs (B) in response to varying concentrations of P5091 at 3d, n = 5. (C) qPCR results of the mRNA levels of *IL-1b*, *Tnf-a*, and *IL-6* in different groups of BMDMs treated P5091, n = 3. (D) ELISA results of IL-1β, IL-6, and TNF-α secretion level in different groups of BMDMs treated with P5091, n = 4. (E) Representative images of ALP/ARS staining of BMSCs co-cultured with BMDMs. (F) WB results and quantification of the protein level of ALP, RUNX2, and COL1α1 in BMSCs co-cultured with BMDMs, n = 3. Y-BMSCs, young BMSCs; A-BMSCs, aged BMSCs; shNC, BMDMs transfected with shNC lentivirus; shUsp7, BMDMs transfected with shUsp7 lentivirus; CON, control BMDMs; SEN, senescence BMDMs. Data are shown as the mean ± SD; ns, not significant, **P* < 0.05, ***P* < 0.01, ****P* < 0.001, *****P* < 0.0001.

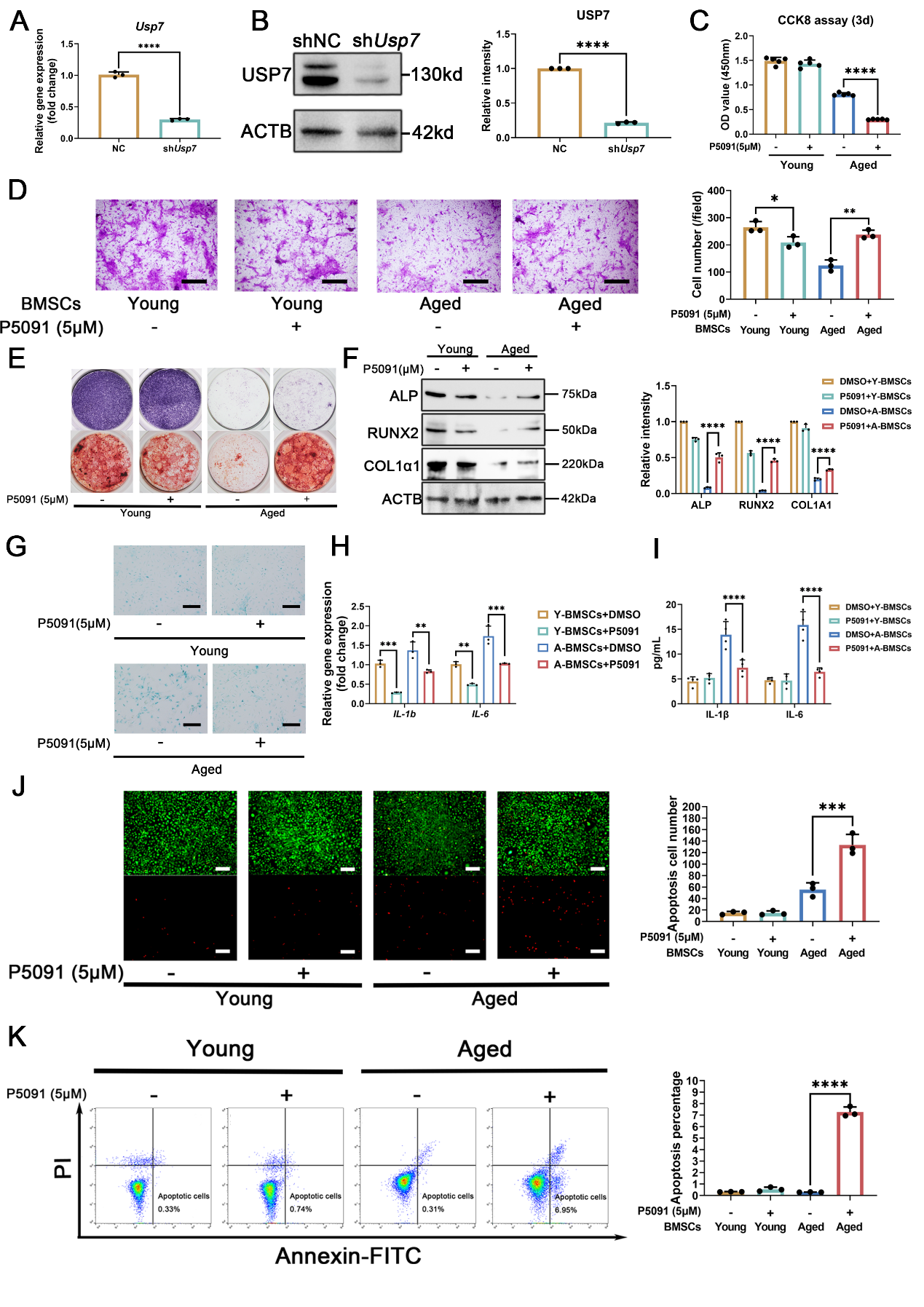

**Appendix Figure 5.** Generation of USP7-depleted BMSCs and P5091 promoted osteogenesis and senolysis in aged mice. (A) qPCR showing downregulated mRNA level of USP7 in BMSCs transfected with sh*Usp7*, n = 3. (B) WB analysis showing of downregulated protein level of USP7 in BMSCs transfected with sh*Usp7*, n = 3. (C) CCK8 assay determining decreased cell viability in aged BMSCs treated with P5091 at 3d, n = 5. (D) Transwell assay indicating that P5091 promoted cell migration in aged BMSCs, n = 3. (E) Representative images of ALP/ARS staining showing that P5091 altered osteogenic capacity in young and aged BMSCs. (F) WB results and quantification determining that altered protein level of ALP, RUNX2, and COL1α1 in young and aged BMSCs treated with P5091 at 7d, n = 3. (G) Representative images of SA-β-gal staining of different groups of BMSCs with P5091, scale bar, 50μM. (H) qPCR results showing that mRNA level of *IL-1b* and *IL-6* in different groups of BMSCs with P5091 at 3d, n = 3. (I) ELISA results showing that secretion level of IL-1β and IL-6 in different groups of BMSCs with P5091 at 3d, n = 4. (J) Representative images and quantitative analysis of live/dead staining of BMSCs after treated with P5091s for 3d, green color indicates live cells, red color indicates dead cells, scale bar, 150μm, n = 3. (K) Flow cytometry and quantitative analysis for apoptosis in BMSCs after treated with P5091 for 3d, n = 3. shNC, BMSCs transfected with NC lentivirus; sh*Usp7*, BMSCs transfected with shUsp7 lentivirus; young, young BMSCs; aged, aged BMSCs. Data are shown as the mean ± SD; ns, not significant, **P* < 0.05, ***P* < 0.01, ****P* < 0.001, *****P* < 0.0001.

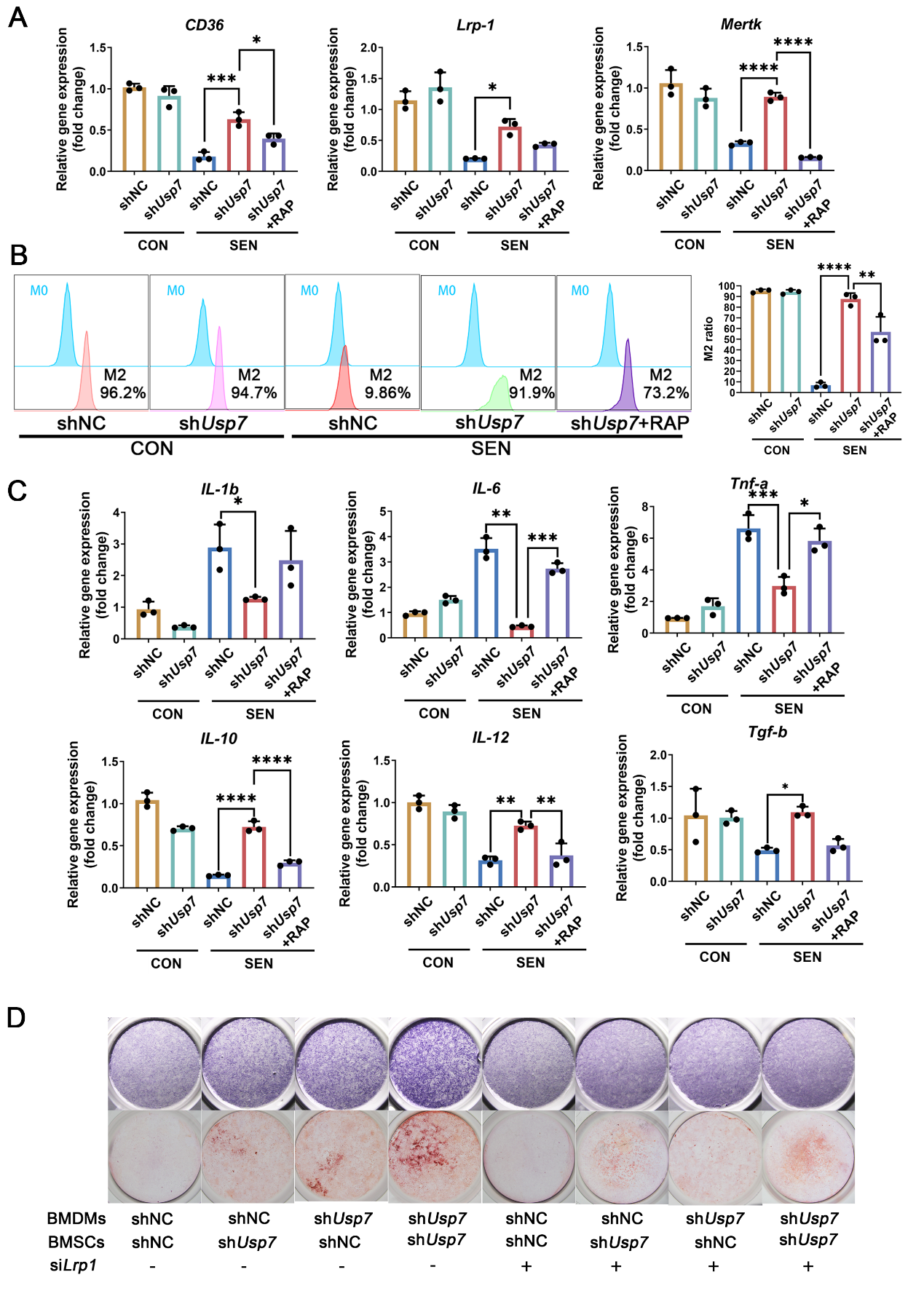

**Appendix Figure 6.** USP7 depletion improved pro-osteogenic capacity and inhibited inflammation in senescent BMDMs engulfing apoptotic cells. (A) qPCR results showing mRNA level of efferocytosis-related genes (*CD36, Lrp1,* and *Mertk*) in BMDMs, n = 3. (B) Flow cytometry showing M2 polarization in different groups of BMDMs that engulfed apoptotic Jurkat cells, n = 3. (C) qPCR results showing mRNA level of pro-inflammatory-related genes (IL-1b, IL-6, and *Tnf-a*) and anti-inflammatory-related genes (*IL-10, IL-12,* and *Tgf-b*) in different groups of BMDMs, n = 3. (D) Representative images of ALP/ARS staining of BMSCs co-cultured BMDMs directly. shNC, BMDMs transfected with NC lentivirus; shUsp7, BMDMs transfected with shUsp7 lentivirus; CON, control BMDMs; SEN, senescence BMDMs; RAP, receptor-associated-protein. Data are shown as the mean ± SD; ns, not significant, **P* < 0.05, ***P* < 0.01, ****P* < 0.001, *****P* < 0.0001.

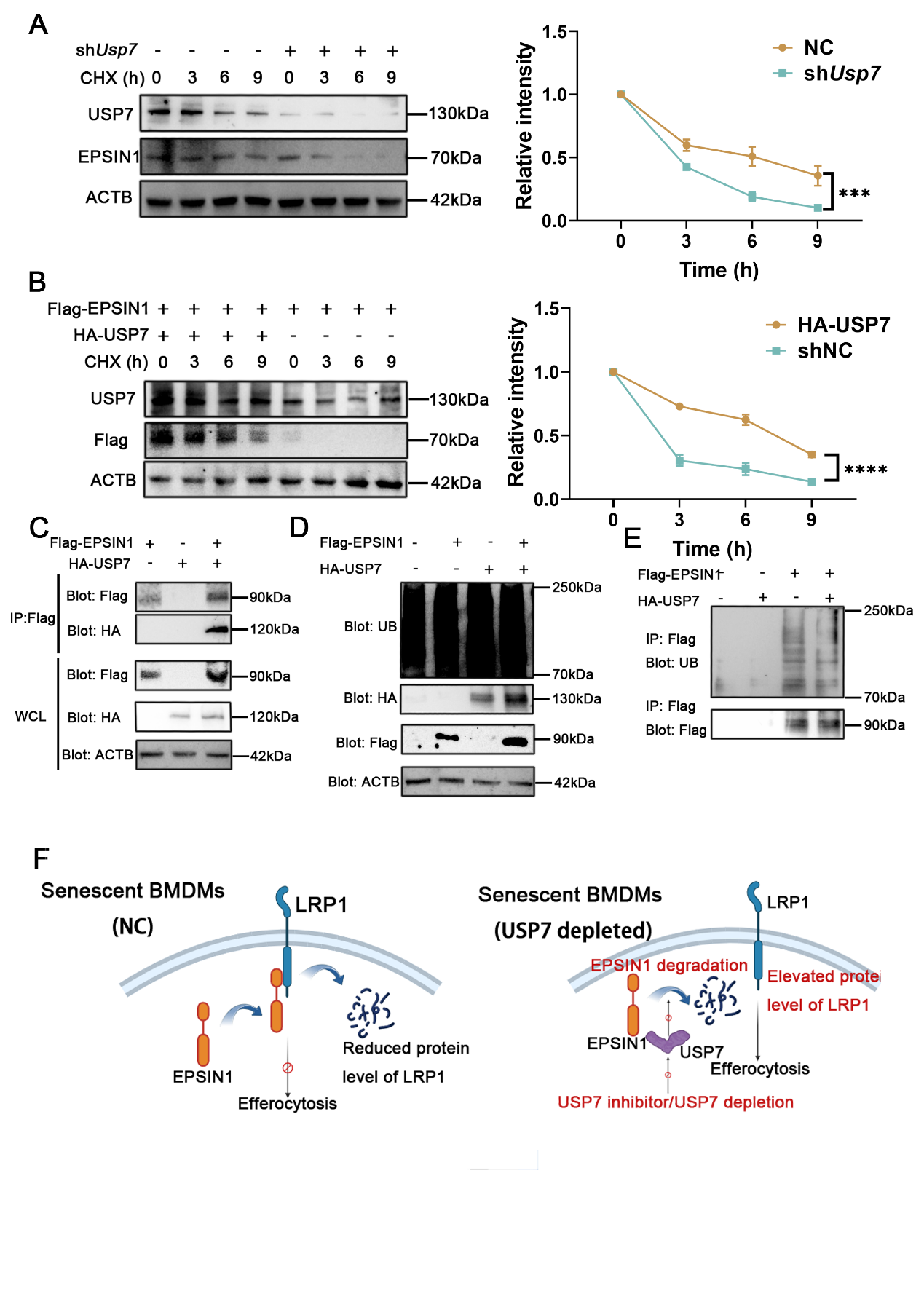

**Appendix Figure 7.** USP7 regulated LRP1-mediated efferocytosis by deubiquitinating EPSIN1. (A) WB results showing the time-dependent expression of EPSIN1 in USP7-depleted BMDMs, n = 3. (B) WB results showing the time-dependent expression of EPSIN1 in BMDMs transfected with HA-USP7, n = 3. (C) Co-IP showing that the interaction between Flag-EPINS1 and HA-USP7 in different groups of BMDMs. (D) WB results showing the overall ubiquitous level in different groups of BMDMs. (E) Co-IP indicating that HA-USP7 over expression downregulated ubiquitous level of EPSIN1. (F) Schematic diagram showing that USP7 regulated LRP1-mediated efferocytosis by deubiquitinating EPSIN1. CHX, cycloheximide; IP, immunoprecipitation; WCL, whole cell lysate. Data are shown as the mean ± SD; ns, not significant, ****P* < 0.001, *****P* < 0.0001.

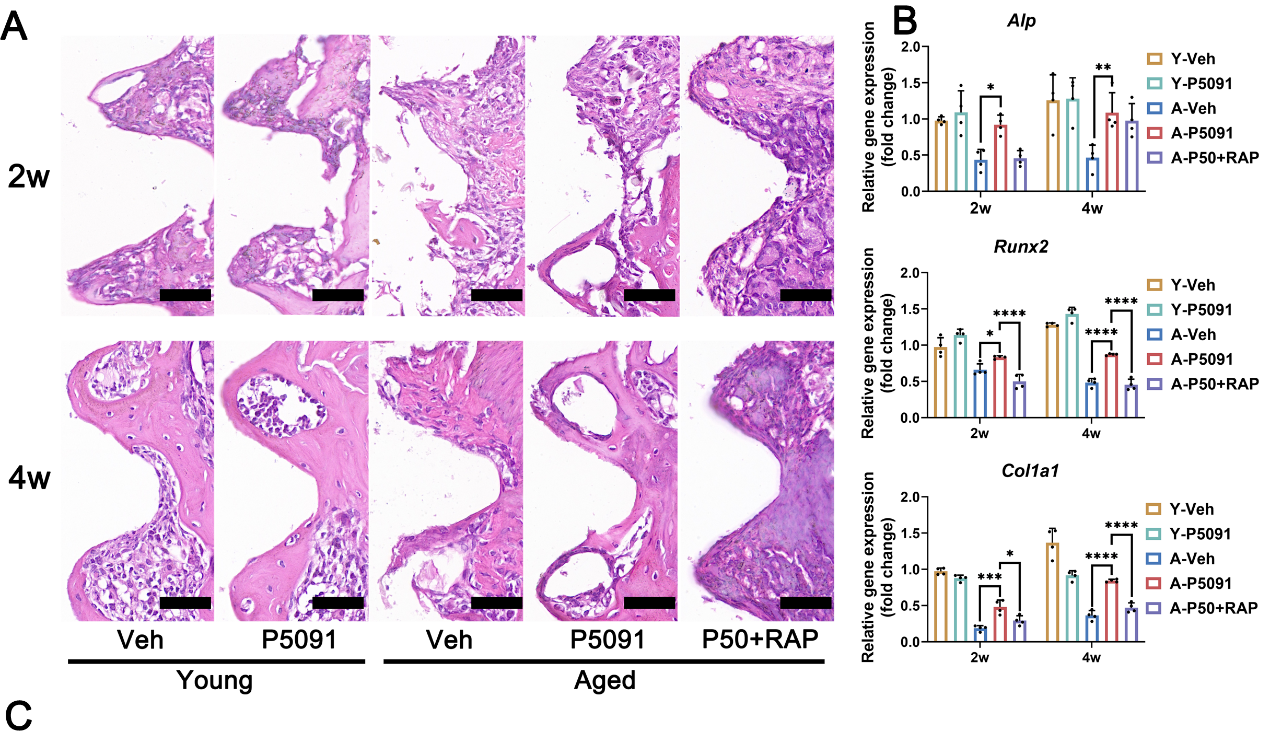

**Appendix Figure 8.** P5091 promoted osteogenesis in peri-implant bone tissue of senile osteoporotic mice. (A) HE staining of peri-implant bone tissue at 2-week and 4-week in different groups of mice. (B) qPCR results showing that mRNA level of *Alp, Runx2,* and *Col1a1* in peri-implant bone tissue of different groups of mice, n = 3. Veh, vehicle; RAP, receptor-associated-protein; Young, young mice; Aged, senile osteoporotic mice. Data are shown as the mean ± SD; ns, not significant, **P* < 0.05, ***P* < 0.01, ****P* < 0.001, *****P* < 0.0001.

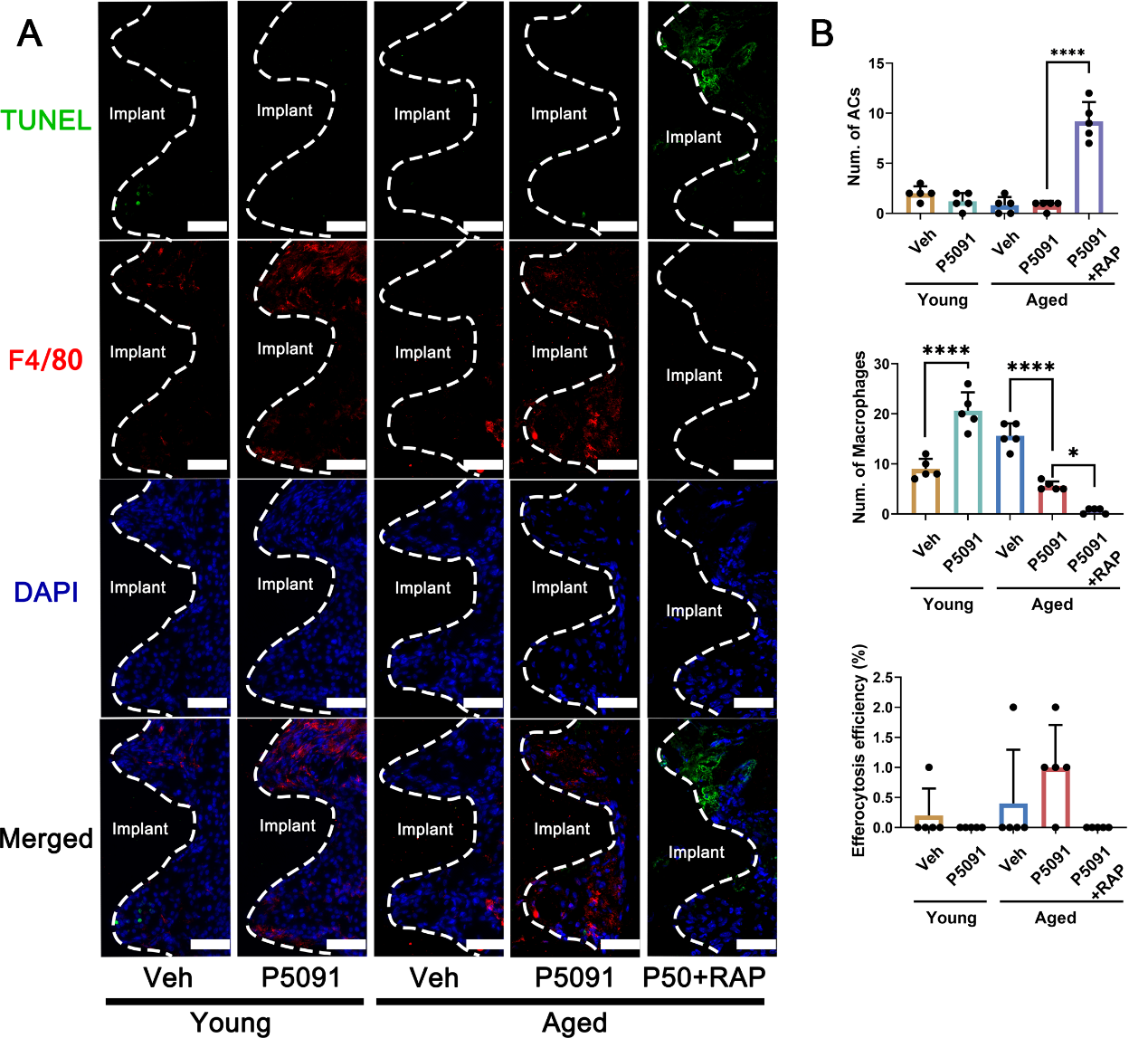

**Appendix Figure 9.** P5091 did not alter efferocytosis at 4-week of osseointegration. (A) Representative immunofluorescence images of macrophages and apoptotic cells around the implants, scale bar, 60μM. Veh, vehicle; RAP, receptor-associated-protein; Young, young mice; Aged, senile osteoporotic mice. Data are shown as the mean ± SD; ns, not significant, **P* < 0.05, ***P* < 0.01, ****P* < 0.001, *****P* < 0.0001.

**Appendix Table 1.** Primers sequences used in this research.

| **Targets (mouse)** | **Primer Sequence** |
| --- | --- |
| *β-actin F* | CATCCGTAAAGACCTCTAGCCAAC |
| *β-actin R* | ATGGAGCCACCGATCCACA |
| *Alp F* | AACCCAGACACAAGCATTC |
| *Alp R* | GCCTTTGAGGTTTTTGGTCA |
| *Runx2 F* | GGTACTTCGTCAGCATCCTATCAG |
| *Runx2 R* | GCTTCCGTCAGCGTCAACAC |
| *Col1α1 F* | GCTTCACCTACAGCACCCTTGT |
| *Col1α1 R* | TGACTGTCTTGCCCCAAGTTC |
| *Il-1β F* | ACCCCAAAAGATGAAGGGCT |
| *Il-1β R* | GATACTGCCTGCCTGAAGCTCT |
| *Il-6 F* | TAGTCCTTCCTACCCCAATTTCC |
| *Il-6 R* | TTGGTCCTTAGCCACTCCTTC |
| *Tnf-a F* | CCCTCACACTCAGATCATCTTCT |
| *Tnf-a R* | GCTACGACGTGGGCTACAG |
| *p53 F* | CTCTCCCCCGCAAAAGAAAAA |
| *p53 R* | CGGAACATCTCGAAGCGTTTA |
| *CD36 F* | ATGGGCTGTGATCGGAACTG |
| *CD36 R* | GTCTTCCCAATAAGCATGTCTCC |
| *Lrp1 F* | ACTATGGATGCCCCTAAAACTTG |
| *Lrp1 R* | GCAATCTCTTTCACCGTCACA |
| *Mertk F* | CAGGGCCTTTACCAGGGAGA |
| *Mertk R* | TGTGTGCTGGATGTGATCTTC |
| *IL-10 F* | GCTCTTACTGACTGGCATGAG |
| *IL-10 R* | CGCAGCTCTAGGAGCATGTG |
| *Tgf-b F* | CTCCCGTGGCTTCTAGTGC |
| *Tgf-b R* | GCCTTAGTTTGGACAGGATCTG |
| *IL-12 F* | CCTGGCTCTTGCTTGCCTT |
| *IL-12 R* | GGTCTTGTGTGATGTTGCTCA |

**Appendix Table 2.** Antibodies used in this research.

| **Antibody** | **Source** | **Host** | **Dilution** |
| --- | --- | --- | --- |
| p16INK4a | Abcam, ab211542 | Rabbit | 1:100 |
| NLRP3 | Adipogen, AB_2490202 | Mouse | 1:1000 |
| cleaved IL-1β | Cell Signaling Technology, Asp116 | Rabbit | 1:1000 |
| cleaved caspase-1(p20) | Adipogen, AG-20B-0042-C100 | Mouse | 1:1000 |
| GSDMD  GSDMD-NT | Huabio, HA601046 | Mouse | 1:1000 |
| β-actin | Huabio, EM21002 | Mouse | 1:1000 |
| ALP | Huabio, ET1609-68 | Rabbit | 1:500 |
| RUNX2 | Huabio, ET161247 | Rabbit | 1:500 |
| COL1α1 | SAB, 28307 | Rabbit | 1:500 |
| USP7 | ThermoFisher, A300-033A | Rabbit | 1:50 |
| EPSIN1 | Abcam, ab75879 | Rabbit | 1:100 |
| LRP1 | Abcam, ab92544 | Rabbit | 1:100 |
| Rabbit Anti-HA | Abcam, ab9110 | Rabbit | 1:100 |
| Rabbit Anti-Flag | Abcam, ab1162 | Rabbit | 1:100 |
| Mouse Anti-HA | Abmart, M20003 | Mouse | 1:100 |
| Mouse Anti-Flag | Abmart, M20008 | Mouse | 1:100 |
| HRP Goat Anti-Rabbit IgG (H+L) | Huabio, HA710308 | Goat | 1:10000 |
| HRP Goat Anti-Mouse IgG (H+L) | Huabio, HA710142 | Goat | 1:10000 |
| F4/80 | Abcam, ab6640 | Mouse | 1:100 |
